# Higher-order epistasis drives evolutionary unpredictability toward novel antibiotic resistance

**DOI:** 10.1101/2025.07.08.663783

**Authors:** Ilona Kamila Gaszek, Muhammed Sadik Yildiz, Devin Meng, Jose Alberto de la Paz, Sophia Maria Alvarez, Faruck Morcos, Milo Lin, Erdal Toprak

**Affiliations:** Department of Pharmacology, The University of Texas Southwestern Medical Center, Dallas, TX 75390, USA; Lyda Hill Department of Bioinformatics, The University of Texas Southwestern Medical Center, Dallas, TX 75390, USA; Department of Biological Sciences, University of Texas at Dallas, Richardson, TX 75080, United States; Department of Bioengineering, University of Texas at Dallas, Richardson, TX 75080, United States; Department of Physics, University of Texas at Dallas, Richardson, TX 75080, United States; Center for Systems Biology, University of Texas at Dallas, Richardson, TX 75080, United States; Department of Biophysics, The University of Texas Southwestern Medical Center, Dallas, TX 75390, USA; Green Center for Systems Biology, The University of Texas Southwestern Medical Center, Dallas, TX 75390, USA

## Abstract

The rapid evolution of extended-spectrum β-lactamases (ESBLs) represents a global health threat, undermining the efficacy of β-lactams, the most extensively used antibiotic class. To elucidate the evolutionary dynamics underlying β-lactam resistance, we constructed a comprehensive combinatorial mutant library comprising all 55,296 possible TEM-1 β-lactamase variants integrating 18 clinically observed mutations across 13 key residues. Over eight million empirical fitness measurements were obtained under selection pressure with both a native antibiotic substrate (ampicillin) and a novel antibiotic (aztreonam), establishing the largest experimentally determined fitness landscape for antibiotic resistance to date. Through graph-theoretic and epistatic analyses, we discovered that selection with ampicillin resulted in weak epistasis, with mutants rarely surpassing the fitness of the wild-type enzyme. Conversely, aztreonam selection elicited extensive higher-order epistasis, generating a rugged fitness landscape characterized by increased phenotypic unpredictability. Interpretable machine-learning analyses identified context-dependent epistatic interactions necessary for achieving high-level aztreonam resistance. Further evolutionary statistical analyses, including direct coupling analysis and latent generative landscapes, showed that top-performing TEM-1 variants consistently adhered to conserved epistatic patterns found in naturally occurring β-lactamases. Our findings demonstrate that higher-order epistasis critically shapes fitness landscape ruggedness when enzymes adapt to novel substrates, whereas adaptations to native substrates exhibit predictably smoother landscapes. This integrated experimental and computational framework provides a foundation for predictive evolutionary pharmacology, enabling assessments of newly developed β-lactams or emerging β-lactamase variants for their potential contribution to ESBL evolution. Importantly, incorporating graph-theoretically informed evolutionary constraints can strategically disrupt evolutionary pathways, presenting a viable approach to mitigate the rise of antibiotic resistance.

## INTRODUCTION

One of the central questions in protein evolution is how mutational pathways enable access to novel functionalities and to what extent chance versus necessity governs this dynamical process. These evolutionary dynamics can, in principle, be conceptualized as trajectories across a fitness landscape, where fitness is mapped as a function of genotype coordinates^1,2^. While extensive theoretical work has explored evolutionary accessibility and reproducibility in terms of fitness landscape features—such as the number of fitness peaks—these models may not capture key properties of empirical fitness landscapes^3,4^.

Traditionally, local empirical landscapes have been constructed through random mutagenesis around single genotypes^5,6^ (commonly the wild type allele). However, such landscapes are combinatorially incomplete, lacking the full set of possible mutational combinations needed to evaluate viable evolutionary trajectories. Recent work has demonstrated combinatorially complete libraries at three^7^ and four^8^ amino acid loci, revealing that evolutionary dynamics can be predictable despite the presence of many fitness peaks. Additionally, combinatorial studies have identified key biophysical constraints driving protein coevolution by quantifying epistatic interactions, particularly at protein–protein interfaces^9^.

Nonetheless, the number of mutated loci in these studies remains insufficient to capture the full spectrum of non-additive fitness effects—known as epistasis—that shape the evolvable structure of the fitness landscape^10^. Current methods do not yet enable the construction of combinatorially complete libraries at the scale of entire proteins, limiting our ability to resolve the global topology of these landscapes. Critically, we also lack protein-wide, combinatorially complete empirical fitness landscapes measured under novel selection conditions that drive the emergence of new functionality.

A pressing example of the evolution of novel protein function is the emergence of antibiotic resistance— particularly the ongoing expansion of extended-spectrum β-lactamases (ESBLs), which render many novel β-lactam antibiotics ineffective. This is a critical public health problem, as antibiotics represent one of the most significant medical advances in human history, enabling not only the treatment of bacterial infections but also essential interventions such as surgeries, organ transplants, and cancer therapies. Among these antibiotics, β-lactams account for nearly 70% of all clinically used injectable antibiotics ^11^.

A major mechanism of β-lactam resistance is the acquisition of β-lactamase genes, often carried on mobile genetic elements^12^. β-lactamases inactivate β-lactam antibiotics by hydrolyzing the β-lactam ring, rendering the drug molecules ineffective (Figure 1A). This form of resistance is particularly concerning because the genes encoding β-lactamases can spread rapidly via horizontal gene transfer. Furthermore, these genes frequently accumulate additional mutations (Figure 1B), giving rise to a broad array of ESBLs capable of degrading newer β-lactam compounds. As a result, ESBL-producing bacteria have been classified as an urgent public health threat by the CDC.

**Figure 1:**
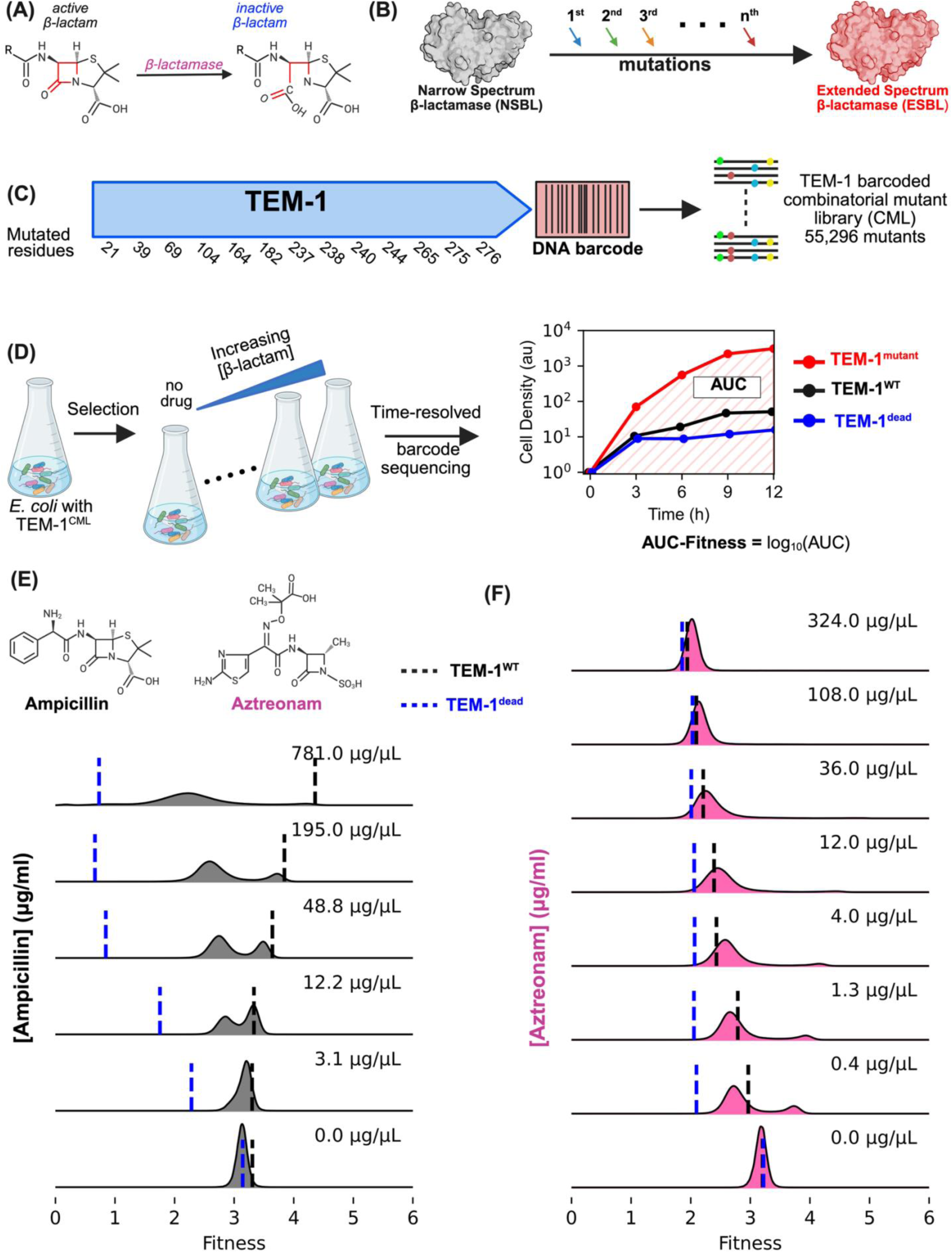
Experimental strategy and global fitness profiling of a combinatorially complete TEM-1 β-lactamase mutant library. (A) β-Lactamases hydrolyze the β-lactam ring, converting an active antibiotic (left) into an inactive product (right), thereby conferring resistance. (B) Clinical evolution from a narrow-spectrum β-lactamase (NSBL, grey surface representation) to an extended-spectrum β-lactamase (ESBL, red) proceeds through the sequential accumulation of point mutations (colored arrows). (C) Schematic of the TEM-1 reading frame showing the 13 positions at which 18 clinically prevalent substitutions were introduced. Exhaustive combinatorial mutagenesis plus random 15-nt DNA barcoding (∼300 barcodes per genotype) yielded a barcoded combinatorial mutant library (TEM-1^CML^) containing all 55,296 possible genotypes. (D) Pooled selection workflow. Escherichia coli carrying TEM-1^CML^ were grown in parallel flasks with increasing concentrations of a β-lactam. Barcode sequencing every 3 h enabled time-resolved tracking of each variant’s abundance. Fitness at each specific dose of β-lactam were quantified as AUC-fitness = log₁₀(AUC), where AUC is the area under the growth curve of each mutant normalized to its cell count at the start of the experiment (right). Curves illustrate a highly resistant mutant (TEM-1^mutant^, red), wild-type TEM-1 (TEM-1^WT^, black) and a catalytically inactive double mutant (TEM-1^dead^, blue). (E) Kernel-density distributions of AUC-fitness scores for the entire library exposed to six concentrations of the native substrate ampicillin (0–781 µg/mL, grey). Black dashed lines mark the median fitness of TEM-1^WT^ and blue dashed lines mark TEM-1^dead^. (F) Equivalent analysis for eight concentrations of the novel monobactam aztreonam (0–324 µg/mL, pink). Compared with ampicillin, aztreonam selection reveals a much broader, high-fitness tail, indicating extensive latent potential for extended-spectrum resistance.

Penicillin and several other β-lactam compounds were originally isolated from microorganisms. By necessity, bacteria that produce β-lactams must evolve resistance mechanisms to protect themselves from the toxic effects of the compounds they synthesize. Likewise, microbial species exposed to β-lactams must develop resistance to survive. As a result, resistance genes—such as those encoding β-lactamases—are common in environmental bacteria and are often disseminated via horizontal gene transfer. To maintain a competitive advantage for access to nutrients and space, β-lactam-producing organisms must also preserve the antibacterial efficacy of their compounds as competitors evolve resistance. Over time, this arms race has driven the diversification of β-lactam molecules, likely in response to the emergence of more effective β-lactamases or ESBLs. Notably, in response to β-lactamase-producing rivals, some bacteria also secrete β-lactamase inhibitors such as clavulanic acid^11^.

This microbial arms race mirrors our own efforts to counter ESBL-producing pathogens by modifying existing β-lactams or developing novel compounds. However, unlike bacteria, our interventions cannot rely on natural selection acting across massive populations. Clinicians must treat every infection— regardless of resistance status—without contributing further to the antibiotic resistance crisis. Therefore, it is essential to quantitatively assess the evolvability of clinically important β-lactamases and forecast the evolutionary trajectories leading to ESBL emergence. A common scientific goal in the infectious disease field is to identify evolutionary “roadblocks” that preemptively delay or prevent resistance development and, ideally, to design β-lactams resistant to evolutionary escape.

We constructed a combinatorially complete fitness landscape encompassing all possible combinations of 18 clinically relevant mutations across 13 residues in the TEM-1 β-lactamase, resulting in 55,296 TEM-1 variants carrying up to 13 mutations. These variants were evaluated under selection by both a β-lactam that TEM-1 efficiently hydrolyzes (ampicillin) and a novel β-lactam to which TEM-1 has negligible activity (aztreonam), across varying concentrations, yielding over 8 million fitness measurements. We show that selection with aztreonam leads to a proliferation of fitness peaks driven by high-order epistasis, resulting in increased sensitivity of the landscape to selection pressure and diminished predictability of mutant fitness. In contrast, under ampicillin selection, epistasis among the tested mutations is weaker, and the fitness landscape remains more predictable. We further demonstrate that both simple epistasis models and machine learning approaches can accurately predict mutant fitness and identify cooperative mutational units that mediate resistance to ampicillin and aztreonam—units also reflected in natural sequence variation. Specifically, we highlight the utility of SHAP scores in interpreting machine learning model outcomes and visualizing the fitness contribution of each mutation when mapping genotypes of the 55,296 TEM-1 variants to their corresponding fitness values. Despite this predictive power, the resulting fitness landscape features a broad array of peaks, complicating optimization for new functionality and highlighting the dominant role of epistasis in shaping evolutionary outcomes.

Finally, our evolutionary statistical analyses using direct coupling analysis (DCA) and latent generative landscapes (LGLs) further demonstrated that high-performing TEM-1 variants consistently adhered to conserved epistatic constraints found in natural β-lactamases supporting the hypothesis that retaining favorable epistatic interactions is essential for preserving TEM-1 function while acquiring resistance to new β-lactams.

## RESULTS

### The combinatorial mutant library of TEM-1 (TEM-1^CML^) exhibits elevated extended-spectrum β-lactamase (ESBL) activity against multiple β-lactam antibiotics

We studied the evolution of a serine β-lactamase into an extended-spectrum β-lactamase (ESBL) by engineering a combinatorial mutant library (CML) of the TEM-1 β-lactamase—a clinically prevalent resistance gene with over 200 reported clinical variants (Figure 1C). This library, referred to as TEM-1^CML^, comprises all 55,296 possible mutants, representing every combination of 18 mutations across 13 targeted residues of TEM-1. These residues and mutations were selected based on their frequent occurrence in clinical TEM-1 variants (Figure S1).

All of the mutated residues, except one (L21), are located within the protein-coding region. L21 resides in the signal sequence of TEM-1, and mutations at this position may alter β-lactamase activity by affecting the periplasmic localization and abundance of TEM-1 variants, rather than through structural changes in the mature protein that could influence substrate-specific activity. Of note, we generated the **L21P** mutation due to a mistake when generating **L21F**, a common mutation found in clinical isolates and known to affect protein localization and thus overall activity. We chose to retain L21P in our analysis to assess its potential impact on fitness.

The **TEM-1^CML^** library was synthesized as a pooled construct, with each variant tagged by numerous randomized DNA barcodes to enable high-resolution, cost-effective, sequencing-based fitness measurements (Figure 1C–D). The library was transformed into an antibiotic-sensitive *Escherichia coli* strain (*E. coli*, NEB10-beta). As controls, *E. coli* cells transformed with a catalytically inactive mutant (**TEM-1^dead^**, carrying the S70A and E166A mutations) were spiked into the library as a reference. In addition, because our library also contained unmutated TEM-1 variants, we designated their fitness as **TEM-1^WT^** in all our analyses and used them as an additional reference. To map barcodes to specific TEM-1 variants, the barcoded library was initially sequenced using long-read PacBio sequencing, generating a reference matchbook that linked each barcode to its corresponding variant. On average, we assigned approximately 300 unique barcodes per variant. All subsequent fitness measurements were performed using 150-nucleotide paired-end Illumina barcode sequencing, providing high resolution at a reduced sequencing cost.

Upon generating the TEM-1^CML^, we screened its collective resistance against a panel of antibiotics from various β-lactam classes, including ampicillin (a natural substrate of TEM-1), cephalosporins, a monobactam, and carbapenems—often considered last-resort antibiotics (Figure S2). As expected, most TEM-1^CML^ mutants did not exhibit improved resistance to ampicillin compared to TEM-1^WT^ (Figure S2), consistent with TEM-1’s already near-optimal intrinsic activity against this compound. Similarly, no enhanced activity was observed against the carbapenems ertapenem, imipenem + cilastatin, or meropenem (Figure S2). Given the limited activity of TEM-1^WT^ against carbapenems, these findings suggest that carbapenem resistant mutants are either absent from the library or extremely rare. In contrast, TEM-1^CML^ demonstrated elevated resistance to all tested cephalosporins—ceftriaxone, cefotaxime, cefepime, and ceftazidime—as well as to aztreonam, a monobactam. Strikingly, the minimum inhibitory concentrations (MICs) for ceftazidime and aztreonam increased by approximately thousand-fold, indicating the presence of a substantial number of highly resistant mutants against these two antibiotics (Figure S2).

### Wild-type TEM-1 exhibits near-optimal activity against ampicillin but shows limited activity against aztreonam

Based on these results, we selected ampicillin (representing the penicillin class for which TEM-1 natively has high activity) and aztreonam (representing the novel monobactam class) for subsequent fitness assays, as they exhibited a large dynamic MIC range—approximately 1,000-fold—between the inactive control (TEM-1^dead^) and the mutant library (TEM-1^CML^). Fitness changes under ampicillin selection were used as a quantitative measure of loss-of-function mutations, while fitness changes under aztreonam selection reflected evolutionary steps toward extended-spectrum β-lactamase activity. Although either ceftazidime (a cephalosporin) or aztreonam could have been used for this purpose, we selected aztreonam because monobactams are relatively less studied in this context, and its chemical structure closely resembles that of ceftazidime—both share an identical R1 side chain containing an aminothiazolyl oxime side chain (Figure 1E).

We quantified the fitness of 55,296 mutants in TEM-1^CML^ under selection with increasing concentrations of either ampicillin or aztreonam using barcode sequencing. Selection experiments were conducted at six concentrations of ampicillin and eight concentrations of aztreonam in triplicates, each including no-drug controls, resulting in approximately 8 million individual fitness measurements. Due to the non-monotonic growth of bacteria under β-lactam selection and the time delay associated with the killing action of β-lactams, we monitored culture growth over a 12-hour period. Samples were collected at 3-hour intervals to measure both cell density (OD_600_) and mutant frequency (*f*). Mutant frequencies were determined by barcode sequencing (Methods). For each mutant, we generated a normalized growth curve, as illustrated in **Figure 1D**, by calculating normalized cell densities using the following formula: 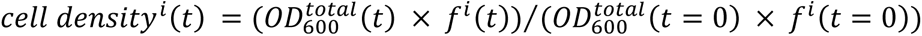, where *i* denotes mutant identity, *t* is time, and *f*^*i*^(*t*) represents the mutant frequency within the library at time *t*.

To quantify mutant performance comprehensively, we defined fitness as the log_10_-transformed area under these normalized growth curves (AUC), hereafter termed AUC-Fitness (Figure 1D). While less conventional than traditional fitness measures such as mutant enrichment after a fixed selection interval or exponential growth rate, AUC-Fitness integrates multiple biologically relevant aspects of growth, including duration of lag phase, growth rate, and final cell density under antibiotic selection. This holistic approach is particularly useful under strong β-lactam selection pressures, where bacterial growth rarely follows simple exponential growth dynamics, thereby limiting the interpretability of classical growth rate metrics. Moreover, the AUC-Fitness metric exhibited excellent reproducibility across biological replicates, typically yielding standard deviations below 10%. Notably, highly resistant mutants typically showed short lag phases, rapid growth rates, and high final cell densities, resulting in significantly greater AUC-Fitness values compared to less resistant mutants, which had longer lag periods, slower growth, and lower final densities (illustrated in Figure 1D).

Importantly, because all mutants are pooled in the same culture, those with high β-lactamase activity can degrade the antibiotics, potentially allowing nearby “cheater” mutants to grow ^13^. However, these cheaters typically start growing later, have longer lag phases, and reach lower final densities because the resistant mutants quickly consume available nutrients and occupy space. Consequently, their AUC values and corresponding AUC-Fitness scores remain low.

As demonstrated in **Figures 1E-F**, the dynamic range of AUC-Fitness measurements vary with selection strength. Under ampicillin selection (**Figure 1E**), the fitness gap between TEM-1^WT^ and TEM-1^dead^ increases with rising ampicillin concentrations, and mutants with fitness values exceeding that of TEM-1^WT^ are rare. In contrast, under aztreonam selection (**Figure 1F**), TEM-1^WT^ exhibits higher fitness than TEM-1^dead^ at low concentrations, but hundreds to thousands of mutants display substantially greater resistance than TEM-1^WT^. At the highest aztreonam concentration, most of the library fails to survive, and the population becomes dominated by approximately 200 highly resistant TEM-1 variants (Figure 1F, **Figure S3**).

### Fitness landscape graphs reveal the impact of selection type and strength on the evolution of extended-spectrum β-lactamase (ESBL) activity

We used graph theory to illustrate the complexity and dynamic structure of the TEM-1 fitness landscape. The concept of a fitness landscape is widely employed in biology to infer evolutionary dynamics and principles from fitness measurements. However, projecting high-dimensional genotype– fitness data onto interpretable, low-dimensional manifolds suitable for visualization remains a significant challenge. Moreover, the large number of mutants (55,296 in our study) can obscure key structural features of the landscape.

A graph-theoretic framework provides a scalable solution for visualizing the fitness landscape, accommodating an arbitrary number of mutations while enabling systematic coarse-graining to extract essential features of the fitness landscape. We developed a graph-based representation of the TEM-1 fitness landscape that preserves key characteristics such as local and global fitness peaks, as well as the mutational pathways that lead to or diverge from these peaks (Figure 2). To construct the graph, each mutant was represented as a node, with edges connecting mutants that differ by a single point mutation. We modeled point-mutation trajectories under a simplified rule: transitions along an edge were permitted only if they led to mutants with equal or higher fitness (i.e., neutral or beneficial mutations). Sometimes multiple such transitions are possible, leading to branching trajectories. To define neutrality, we applied a fitness difference threshold that captures 99% of all pairwise fitness difference amplitudes between edge-connected mutants in the absence of drug-induced selection (Figure 2A). We then applied an iterative coarse-graining process in which all mutants connected by neutral edges were merged into a single node, continuing this process until no further merging was possible. The effective edge weight between any two coarse-grained nodes was computed based on the sum of all fitness differences across edges connecting their constituent members.

**Figure 2:**
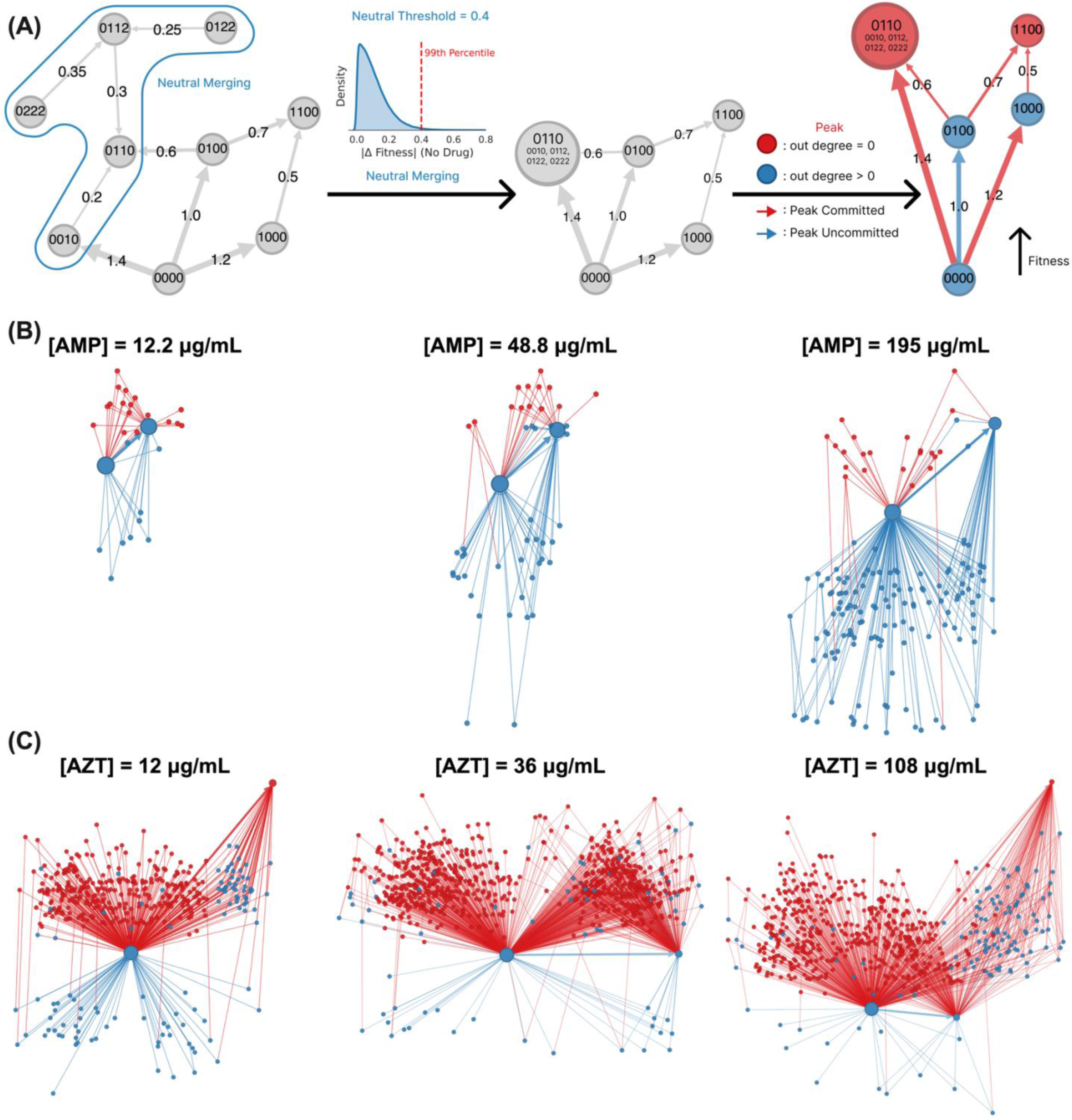
Fitness landscape graphs of TEM-1 under different selection pressures. (A) Algorithm to generate the fitness landscape graph from combinatorial saturation mutagenesis data. Starting from an initial graph in which each node is a mutant and edges connect mutants separated by a point mutation. Nodes connected by edges whose fitness difference fall below the neutral cutoff (dashed line in the density plot of |Δfitness| in the absence of drug) are iteratively merged into supernodes to yield a simplified graph in which peak nodes (red) and connection nodes (blue), with arrows showing committed (red) versus uncommitted (blue) evolutionary flows. (B) The fitness landscape graph under increasing ampicillin ([AMP] = 12.2, 48.8, 195 µg/mL; B) and aztreonam ([AZT] = 12, 36, 108 µg/mL; C) concentrations. Y-axis of graph is proportional to fitness, with fitness determined by mean and max fitness of the mutants in the connection (blue) and peak nodes (red), respectively. Node sizes are monotonically scaled to the number of mutants contained in each node.

In the final graph, all nodes fell into one of two categories. Peak nodes (red in Figure 2A) represent groups of mutants that form standing variation around a local fitness maximum and have no outgoing edges. Connection nodes (blue in Figure 2A) possess at least one outgoing edge and represent intermediate genotypes that can continue to evolve. In this framework, a trajectory originating from a peak node cannot leave it, while a trajectory beginning from a connection node will eventually reach and remain within one of the peak nodes. This simplified transition model implies that all mutational paths ultimately converge on a peak node. A committed transition—a step after which the evolutionary trajectory is guaranteed to reach a specific peak—is represented by red arrows in Figure 2A. In contrast, non-committed transitions, which allow for multiple potential peak outcomes, are shown as blue arrows.

Using this algorithmic mapping, we converted the local fitness measurements from the TEM-1^CML^ dataset (Figure 1E–F) into fitness landscape graphs for each drug and concentration condition (Figures 2B–C for selected conditions; Figure S4 for all concentrations). For **ampicillin**, the resulting graphs display a small number of peak nodes (Figure S4A). Regardless of the starting genotype in the combinatorial library, trajectories consistently converge on a limited set of genotype clusters—each representing standing variation near a local fitness maximum. This structure remains consistent across increasing ampicillin concentrations (Figure 2B), suggesting that the evolutionary dynamics of this well-optimized enzyme are characterized by low fate diversity and robustness to changes in selection pressure.

In contrast, under aztreonam selection, the TEM-1 fitness landscape fragments into hundreds of combined peak nodes (Figure 2C; Figure S4B), indicating a much more diverse array of possible evolutionary outcomes. In this context, the final evolutionary trajectory is highly dependent on the starting genotype. Moreover, due to the stochastic nature of transitions between neutral nodes, even trajectories originating from the same genotype can diverge and reach different fitness peaks. This reflects a highly permissive evolutionary process under aztreonam selection.

Notably, the aztreonam fitness landscape exhibits a strong and non-monotonic dependence on drug concentration. At both low and high aztreonam concentrations, a prominent and well-populated global fitness maximum is clearly distinguishable from other peaks. However, at intermediate concentrations, this global peak disappears as it becomes absorbed into a large connection node, eliminating its influence on evolutionary trajectories (Figure 2C). Selection strength for novel functionality tends to increase the degree of epistasis (as we demonstrate below) and, accordingly, is expected to increase the diversity and history-dependence of evolutionary trajectories. Our simplified graph-theoretic model, applied to the combinatorial mutagenesis data, confirms that selection for novel functionality not only amplifies epistatic interactions but also greatly expands the number of possible evolutionary fates.

### TEM-1^WT^ is multiple mutations away from the optimal fitness peak under aztreonam selection

To study the evolutionary principles shaping the fitness landscape of TEM-1, we focused on measurements conducted at the highest ampicillin concentration (781 μg/mL), where the dynamic range between TEM-1^dead^ and TEM-1^WT^ spanned nearly four orders of magnitude, and at 36 μg/mL aztreonam—a value slightly above the clinical cutoff (32 μg/mL) for aztreonam-resistant E. coli. Although fitness measurements at higher aztreonam concentrations were available to us, we opted against them because most mutants failed to grow under those conditions, resulting in a loss of fitness information.

As shown in Figure 3A, only a small fraction of mutants in TEM-1^CML^ exhibited local fitness values exceeding that of TEM-1^WT^ under ampicillin selection. However, the vast majority still outperformed TEM-1^dead^. This suggests that TEM-1 can tolerate multiple mutations while retaining functionality against ampicillin. Nevertheless, this observation may not be broadly generalizable, as the 18 mutations used in TEM-1^CML^ were selected based on their frequent occurrence in clinical variants and are therefore preselected for functional compatibility. Another possibility is the cooperative effect observed in liquid culture, where highly active variants such as TEM-1^WT^ rapidly hydrolyze ampicillin, lowering the effective drug concentration and indirectly supporting the growth of less active mutants ^13^.

**Figure 3:**
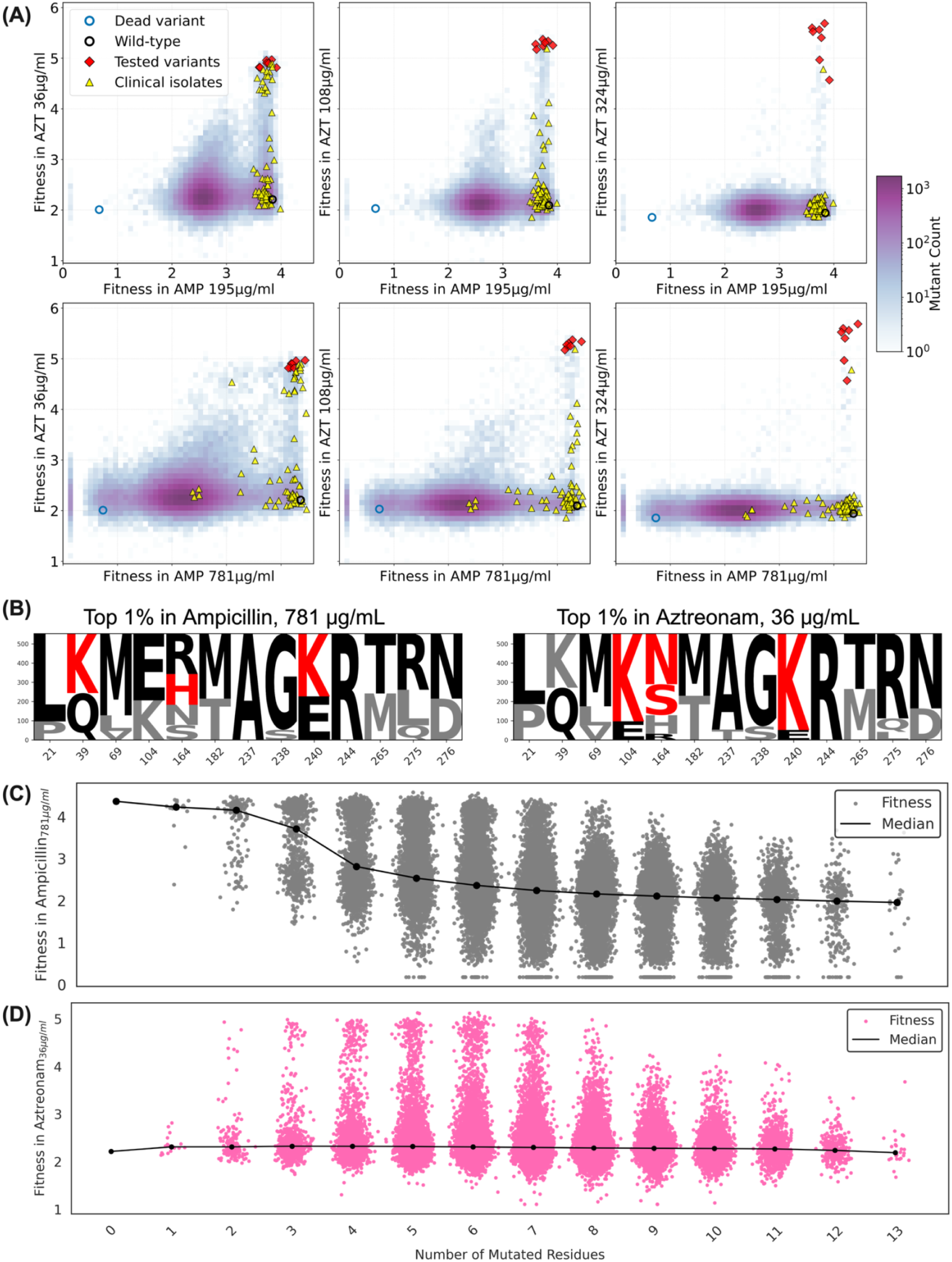
Distinct mutational signatures and epistatic constraints shape the TEM-1 fitness landscape under ampicillin versus aztreonam selection. **(A) Pairwise fitness comparisons at matched antibiotic concentrations.** Each density–scatter plot contrasts fitness (log-transformed area-under-the-growth-curve, AUC-Fitness) in aztreonam (AZT; 36, 108, or 324 µg/mL, y-axes) with fitness in ampicillin (AMP; 195 µg/mL, top row, or 781 µg/mL, bottom row, x-axes). Hex-bin color denotes the number of genotypes (log₁₀ scale). The catalytically inactive reference (TEM-1^dead^, blue open circle), TEM-1^WT^ (black open circle), 7 follow-up variants (red diamonds) and naturally occurring clinical isolates represented in the library (yellow triangles) are overlaid. Very few mutants outperform TEM-1^WT^ in AMP (vertical “plateau” at x ≈ 4), whereas hundreds of variants achieve markedly higher fitness in AZT (upper-right quadrant), especially at elevated AZT concentrations. **(B) Mutational enrichment among the top 1% of variants.** Sequence logos for the 13 interrogated positions were generated from the top 1% of genotypes at 781 µg/mL AMP (left) or 36 µg/mL AZT (right). Letter height is proportional to residue frequency. Wild-type residues are black, enriched substitutions are red. Under AMP selection only Q39K, R164H and E240K show mild enrichment, underscoring the near-optimality of TEM-1 for its native substrate. AZT selection strongly favors E104K, R164H/S/N and E240K, while conserving A237, G238 and R244. **(C, D) Influence of mutational load on resistance.** Fitness of every genotype is plotted against its total number of substitutions for AMP 781 µg/mL (grey, C) or AZT 36 µg/mL (pink, D). Solid black lines denote median fitness. AMP resistance declines monotonically with additional mutations, indicating additive or mildly antagonistic effects. In contrast, AZT resistance displays a broad distribution and peaks in variants harboring four to seven mutations, revealing extensive higher-order positive and negative epistasis. Collectively, these analyses show that the TEM-1 landscape is smooth and mutation-averse under native-substrate pressure but becomes rugged and highly evolvable when challenged with the novel monobactam aztreonam.

In contrast, under aztreonam selection, hundreds of mutants in TEM-1^CML^ displayed high fitness values, highlighting the latent evolutionary potential of TEM-1 to evolve into an extended-spectrum β-lactamase. Strikingly and contrary to our expectations, many aztreonam-resistant mutants also retained high resistance to ampicillin (Figure 3A). This suggests that ESBL evolution may not necessarily involve phenotypic trade-offs for every drug pair and could instead yield variants with strong activity against multiple β-lactam antibiotics simultaneously.

### Epistatic Interactions Govern Mutational Preferences and Fitness Landscapes in TEM-1 β-Lactamase

Next, we analyzed all TEM-1^CML^ mutants by quantifying the frequency of mutations among variants in the top 1% of local fitness under either ampicillin (781 μg/mL) or aztreonam selection (36 μg/mL). As shown in Figure 3B, only three mutations (Q39K, R164H, and E240K, shown in red) at the 13 targeted positions were slightly enriched over the wild-type amino acid (shown in black) under ampicillin selection, underscoring the robust activity of wild-type TEM-1 against this drug. In contrast, under aztreonam selection, variants carrying the E104K, R164H/S/N, and E240K mutations (Figure 3B, red letters) were significantly enriched. Additionally, under both selection conditions, variants retaining the wild-type residues at A237, G238, and R244 were more prevalent. This pattern suggests that E104K, R164H/S/N, and E240K are beneficial for aztreonam resistance, particularly when A237, G238, and R244 remain unmutated. Finally, the L21P mutation was less preferred under both selection conditions, suggesting a fitness cost. Interestingly, however, it was still tolerated in many variants, as also seen among the most resistant variants in Figure S4. To examine how mutational load influences resistance, we plotted AUC-Fitness against the number of mutations per variant across the library. Under ampicillin selection (Figure 3C), the median AUC-Fitness (black line) declined monotonically as the number of mutations increased, indicating a general fitness cost of additional mutations. By contrast, under aztreonam selection (Figure 3D), the median AUC-Fitness remained relatively stable across different mutation numbers, although a wide range of fitness values was observed. Notably, the number of aztreonam-resistant variants and their AUC-Fitness values peaked in mutants carrying **four to seven mutations**, before declining at higher mutational loads. This **non-monotonic trend** suggests the presence of **epistatic interactions**, in which the combined effects of mutations are non-additive and context-dependent.

To assess whether aztreonam resistance could be predicted solely by the **presence of E104K, R164H/S/N, and E240K**, combined with the **absence of mutations at A237, G238, and R244**, we examined all variants matching this mutational profile. This condition failed to fully capture the spectrum of resistant genotypes. This further supports the role of **higher-order positive and negative epistasis** in shaping the aztreonam fitness landscape of TEM-1.

### Higher-order epistasis arises from the suboptimal wild-type TEM-1 sequence for β-lactamase activity against aztreonam

To investigate the degree and extent of epistasis, we first analyzed the effects of single and double mutations on AUC-Fitness under ampicillin (781 μg/mL) or aztreonam (36 μg/mL) selection (Figure 4). Under ampicillin selection, most single mutations were neutral with respect to TEM-1^WT^, although A237T and R244C/S had mild deleterious effects (Figure 4A). When examining all pairwise combinations, several double mutants including those involving G238S, A237T and R244C/S exhibited further reduced AUC-Fitness (Figure 4C). Calculated pairwise epistatic interactions were generally small but included examples of both positive and negative epistasis under ampicillin selection (Figure 4E).

**Figure 4.**
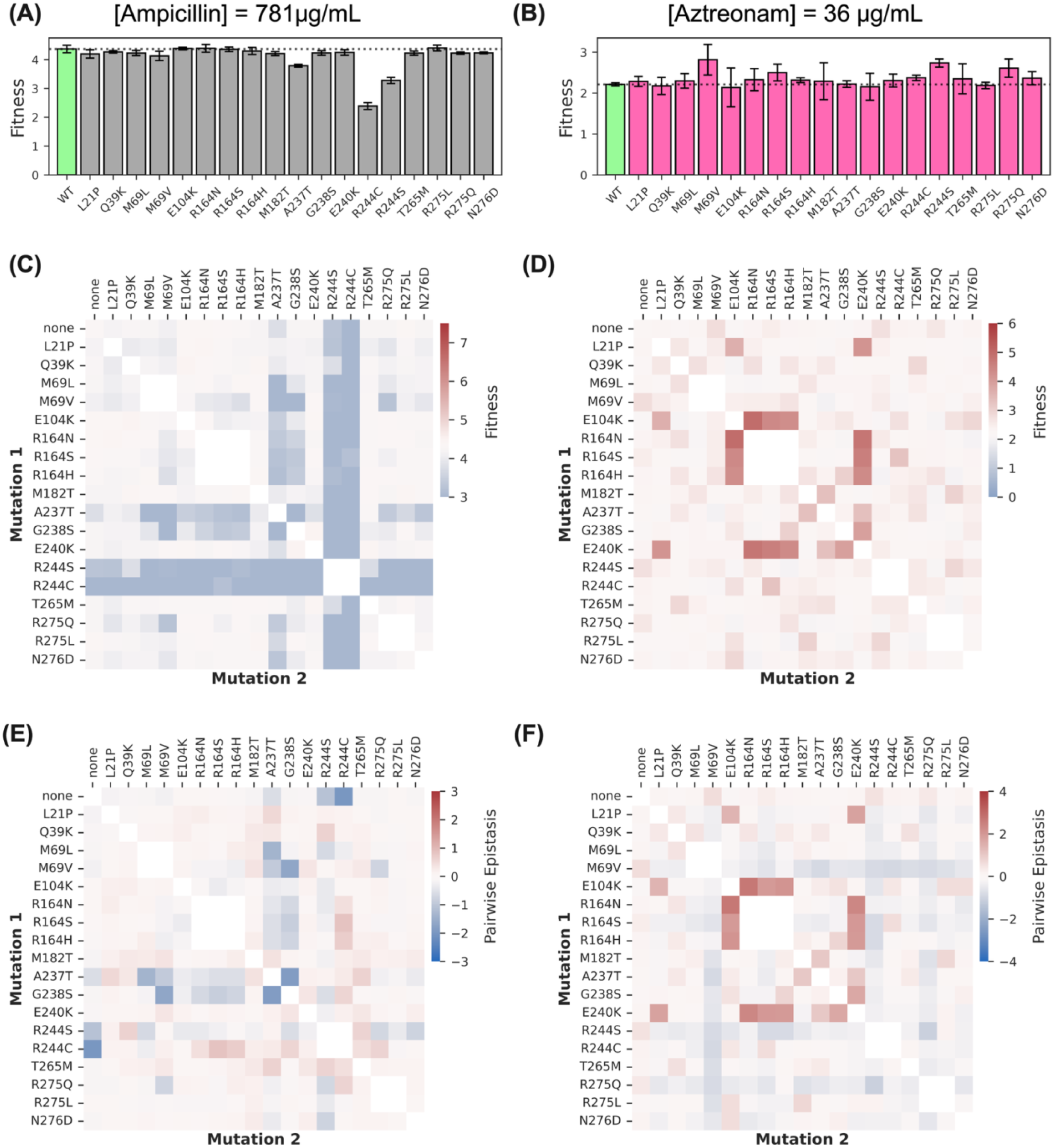
Single- and double-mutant fitness and pair-wise epistasis in the TEM-1^CML^ library. **(A, B)** Fitness of all single mutants (AUC-fitness) after 12 h selection with 781 µg mL^-1^ ampicillin (A) or 36 µg mL^-1^ aztreonam (B). Green bars, wild-type TEM-1 (WT); grey (A) or magenta (B) bars, individual point mutants. Dashed line marks WT fitness. Bars show mean ± SD of three biological replicates. **(C, D)** Heat-maps of mean absolute fitness for every pairwise combination of the 18 mutations at the same drug concentrations. Rows and columns are labelled by the first and second mutation, respectively. The “none” row/column corresponds to single-mutant data and reproduces the values in panels A and B. Color intensity (scale bars at right) reflects AUC-Fitness. **(E, F)** Pair-wise epistasis (ε) for the same double mutants, calculated as ε = F_AB_−F_A_−F_B_+F_WT_, where F is AUC-fitness. Red indicates positive (synergistic) epistasis, blue negative (antagonistic) epistasis; scale bars show ε in AUC-fitness units. Under ampicillin selection (A, C, E) most single mutations are neutral or mildly deleterious and epistatic interactions are small, yielding a relatively smooth landscape. In contrast, aztreonam selection (B, D, F) reveals numerous high-fitness double mutants and widespread, higher-magnitude positive and negative epistasis, particularly among the E104K, R164H/S/N, and E240K mutations, producing a markedly more rugged and unpredictable fitness landscape.

In contrast, under aztreonam selection, many single mutations reduced AUC-Fitness, with only a few mutations such as M69V and R275Q showing mild beneficial effects (Figure 4B). Analysis of double mutants (Figure 4D) and pairwise epistatic interactions (Figure 4F) revealed that combinations involving E104K, R164H/S/N, and E240K often conferred high aztreonam resistance whereas none of the double mutants involving M69V or R275Q displayed high values. These findings indicate widespread positive and negative epistasis particularly among E104K, R164H/S/N, and E240K mutations under aztreonam selection.

To evaluate evolutionary trajectories across the fitness landscape, we analyzed AUC-Fitness values for all combinations of mutations E104K, R164S, and E240K on various genetic backgrounds: TEM-1^WT^ (black), Q39K (blue), A237T (teal), and G238S (orange). These assessments were conducted under selection pressure from ampicillin (Figure S5-A) and aztreonam (Figure S5-B). Under ampicillin selection, all tested mutation combinations demonstrated AUC-Fitness values equal to or slightly below that of the TEM-1^WT^ background, suggesting no significant fitness advantage over the wild-type enzyme. The Q39K background did not alter this trend. However, combinations such as E104K + E240K and E104K + R164S were notably deleterious when placed on either the A237T or G238S backgrounds (Figure S5A).

In contrast, aztreonam selection revealed significant complexity in the fitness landscape due to epistatic interactions (Figure S5B). Starting from the TEM-1^WT^ background, only the R164S mutation provided a slight initial fitness benefit, whereas E104K and E240K alone showed nearly neutral effects. The double mutant (E104K + E240K) exhibited AUC-Fitness values similar to either single mutant. However, when R164S was present, both E104K and E240K mutations displayed strong positive epistasis, becoming significantly beneficial. Moreover, acquiring either E104K or E240K in this context enhanced the fitness even further upon subsequent addition of the third mutation (Figure S5). The Q39K background did not alter these interactions (Figure S5-B, blue). Interestingly, the A237T mutation increased fitness of the E240K single mutant and the E104K + E240K double mutant but negatively impacted the fitness of combinations including E104K + R164S, R164S + E240K, and the triple mutant E104K + R164S + E240K (Figure S5-B, teal). Similarly, the G238S mutation increased the fitness of the E240K single mutant but reduced fitness in combinations involving R164S + E240K and E104K + R164S (Figure S5-B, orange).

These findings highlight the significance of higher-order epistasis in shaping the fitness landscape of TEM-1 under aztreonam selection, demonstrating a more intricate and rugged landscape compared to ampicillin selection. This observation aligns with the broader patterns we have seen in the fitness landscape graphs (Figure 2), underscoring the greater influence of epistatic interactions in ESBL evolution under aztreonam.

### Epistatic Complexity Limits the Predictive Reconstruction of Fitness Landscapes

To evaluate how well partial fitness measurements can reconstruct the full fitness landscape (AUC-Fitness), we used fitness data from low-order mutants (e.g., single and double mutants) to predict the AUC-Fitness of higher-order variants. The completeness of our library allowed us to calculate all epistatic interactions up to 13th order (Figure 5). Then, we applied a linear regression model to predict AUC-Fitness across the entire TEM-1^CML14–16^. As expected from our pairwise epistasis analysis (Figure 4), predictive power was limited under ampicillin (781 μg/mL; Figure 5A, C) or aztreonam (36 μg/mL; Figure 5B, D) selection. Predictive accuracy naturally further improved with the inclusion of second- and higher-order epistatic terms.

**Figure 5.**
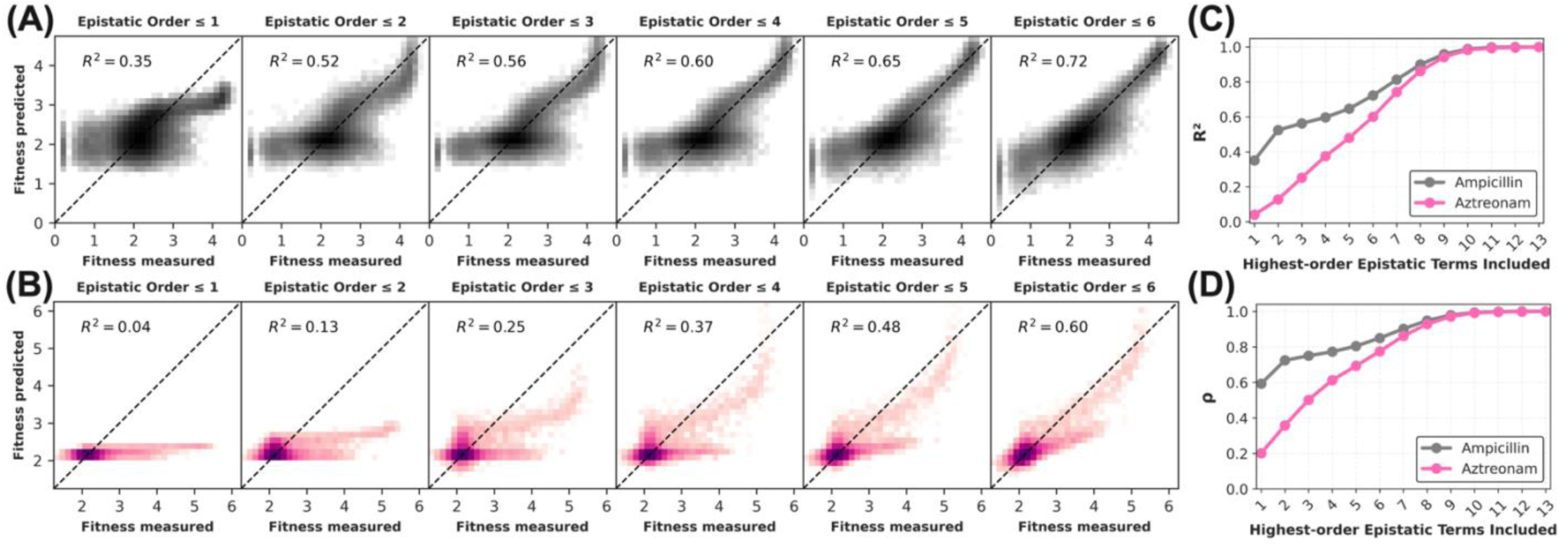
Incorporating higher-order epistasis is essential to accurately reconstruct the TEM-1 fitness landscape, particularly for the novel substrate aztreonam. **(A)** Density plots comparing experimentally measured AUC-Fitness values for all 55,296 TEM-1 variants assayed at 781 µg/mL ampicillin with values predicted by linear models that incrementally include epistatic terms through the indicated maximum order (≤1 additive, ≤2 pairwise, … ≤6). Dashed lines denote perfect prediction and in-panel R² values quantify explained variance. **(B)** Equivalent analysis for aztreonam (36 µg/mL) reveals much poorer predictions when only low-order terms are included, reflecting a rugged, highly epistatic landscape. **(C)** Global coefficient of determination (R²) as higher-order terms are added. For ampicillin (grey), additive + pairwise interactions already explain just over half of the total variance (R² ≈ 0.52), and improvements taper after third-order terms. In stark contrast, aztreonam (pink) does not reach the 0.5 R² threshold until fifth-order terms are incorporated, underscoring the dominance of higher-order epistasis for the novel substrate. **(D)** Rank-order fidelity, expressed as Spearman correlation (ρ), exhibits the same trend: rapid convergence for ampicillin but a steady, higher-order-dependent climb for aztreonam. Together, these results show that the native-substrate landscape is comparatively smooth and predictable, whereas adaptation to a novel substrate exposes extensive higher-order interactions that must be modelled to forecast fitness accurately.

These results suggest that in rugged fitness landscapes where epistasis is widespread, systematically reconstructing the full fitness landscape by incrementally building libraries of single, double, triple, and higher-order mutants is unlikely to be an efficient strategy. Instead, a more practical and scalable approach may involve generating low-order mutants in parallel with random or targeted combinatorial variants, guided by evolutionary statistics derived from naturally occurring and clinically observed TEM-1 sequences.

### Distilling and Visualizing Epistatic Rules Using Machine Learning

We extracted the genetic features of the TEM-1 fitness landscape under ampicillin (781 μg/mL) and aztreonam (36 μg/mL) selection by training a machine learning model to predict the AUC-Fitness of TEM-1 variants carrying any combination of the 18 mutations we studied. Our predictive model links genotype, specifically, the presence or absence of mutations at studied positions to observed fitness values, providing a quantitative framework for genotype-to-phenotype mapping. For this purpose, we employed a Light Gradient Boosting Machine (LightGBM) based regression model, a decision tree– based method capable of capturing complex, non-linear interactions between mutations that influence fitness. Model performance was assessed through learning curve analysis and evaluation on held-out test data to ensure predictive capability (Figure 6A-B).

**Figure 6.**
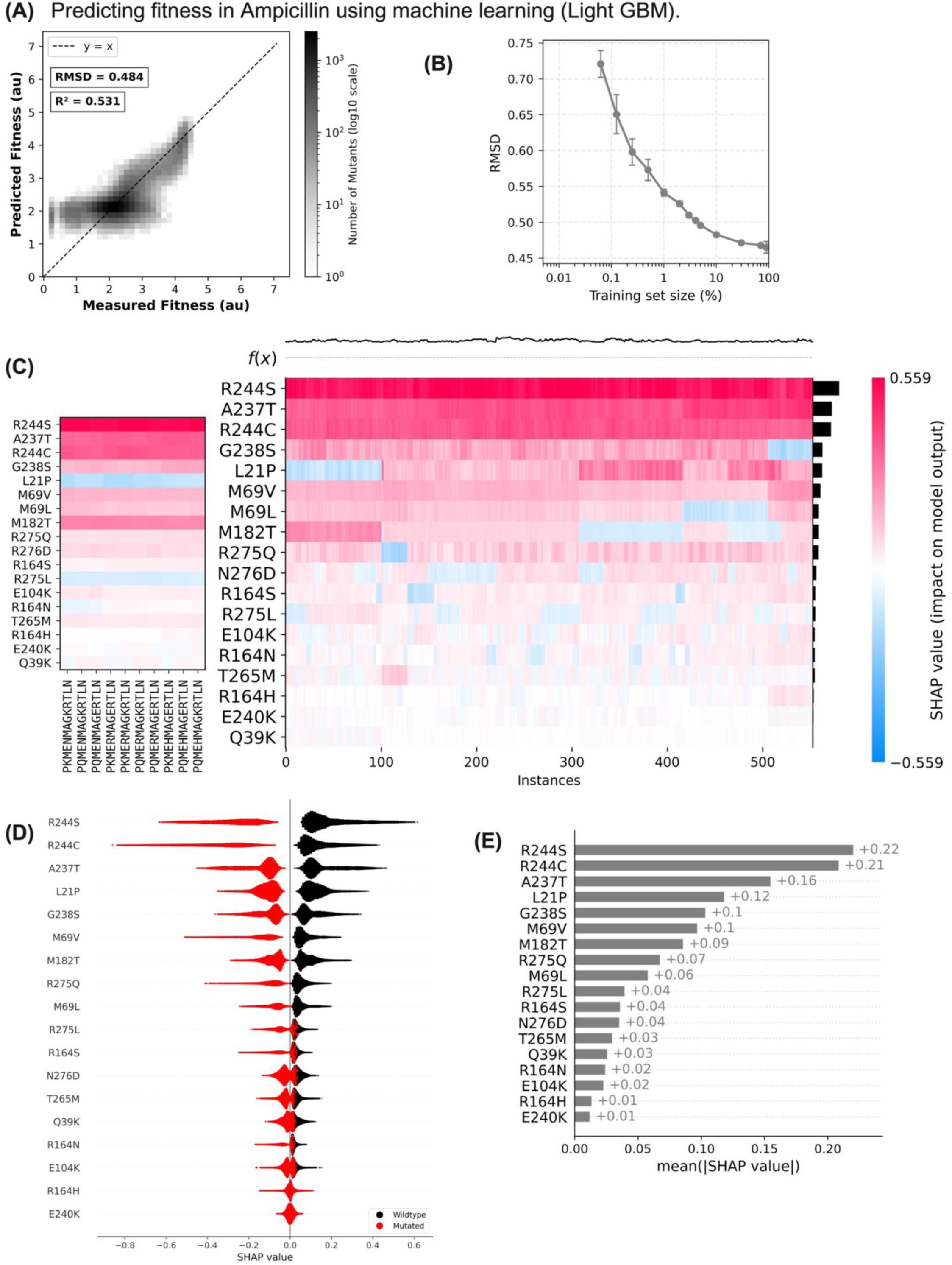
Machine learning prediction and SHAP interpretation of TEM-1 fitness under ampicillin selection. (A) Correlation between experimentally measured and LightGBM predicted AUC-fitness for 55,296 TEM-1 variants. The prediction model was trained using a randomly selected subset comprising 10%; the dashed line indicates a perfect 1:1 correlation. (B) Learning curve analysis showing the root mean square deviation (RMSD) between measured and predicted fitness values (y-axis) as a function of the training set size (x-axis, log scale). Prediction accuracy improves rapidly as training set size increases from 0.03% (∼17 genotypes) up to approximately 1% of the library, beyond which diminishing returns occur. (C) Heatmap depicting SHAP (SHapley Additive exPlanations) values for the 553 most ampicillin-resistant genotypes (∼1% of the library). Columns represent the 18 studied mutations, rows correspond to individual genotypes, and colors indicate each mutation’s contribution to predicted fitness (red: positive; blue: negative; white: neutral). (D) Beeswarm plot summarizing SHAP values across all evaluated genotypes. Each dot represents an individual genotype colored by mutation status (red: mutation present; blue: wild-type residue). Horizontal distribution indicates the magnitude and direction of SHAP contributions. The absence of R244C/S mutations confers the most substantial positive effect, whereas other mutations generally reduce fitness. (E) Mean absolute SHAP values for each position, summarizing the overall magnitude of each mutation’s influence. Mutation at position R244 has the greatest impact, followed by mutations at positions A237, M69, and G238. Mutations Q39K and E240K exhibit minimal contributions under ampicillin selection. Collectively, panels A–E demonstrate that even limited training data enable accurate fitness predictions and highlight key mutations driving ampicillin resistance.

Under ampicillin selection, the model achieved high accuracy in predicting AUC-Fitness across the full TEM-1^CML^ library (Figure 6A–B). The training set size was systematically varied from 0.03% to 99% of the data, and prediction performance was evaluated using RMSD in AUC-Fitness values (Figure 6B). The model demonstrated decent predictive power, which improved rapidly as the training size increased, showing diminishing returns beyond approximately 10% of the data. Figure 6A illustrates the correlation between measured and predicted AUC-Fitness values when 10% of the data was used for training.

When the same modeling approach was applied under aztreonam selection (Figure 7A–B), model performance did not immediately improve with increasing training size—likely because a large portion of variants in TEM-1^CML^ performed poorly under aztreonam selection, and sampling these variants did not aid model learning. However, training performance improved rapidly once the training size exceeded 0.5% of the data. Notably, using just 10% of the data yielded a promising RMSD of 0.288 (Figure 7A), which may be sufficient for qualitatively classifying mutants as “resistant” for clinical purposes.

**Figure 7.**
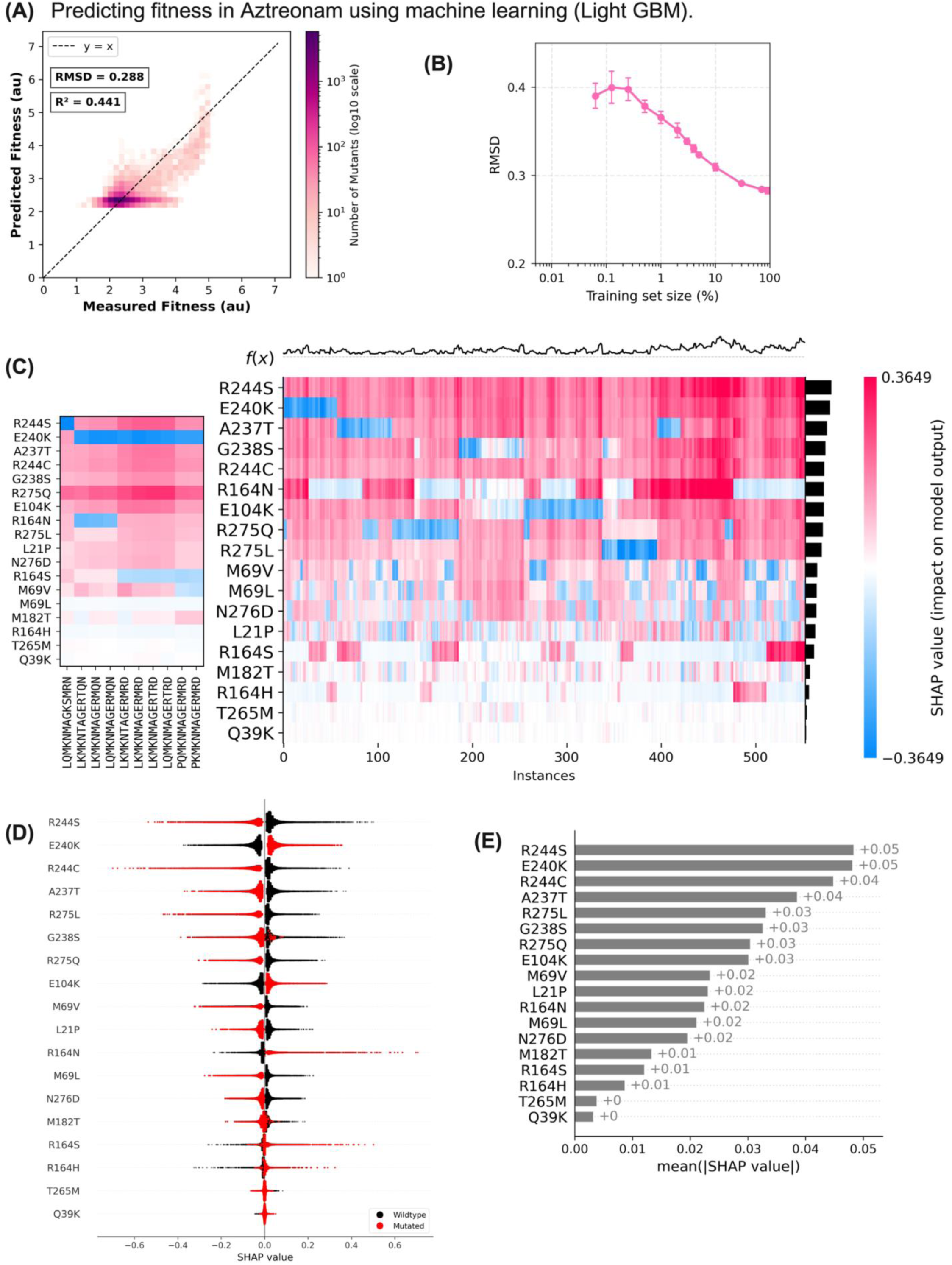
Machine learning prediction and SHAP interpretation of TEM-1 fitness under aztreonam selection. (A) Correlation between experimentally measured and LightGBM predicted AUC-fitness values for 55,296 variants. Predictions utilized a 10% random subset of genotypes for training, consistent with Figure 6A. Increased dispersion (RMSD = 0.426) indicates a more complex fitness landscape under aztreonam selection compared to ampicillin. (B) Learning curve analysis displaying RMSD as a function of training set size. Even when using 99% of available data, the model does not fully converge, reflecting the greater complexity and ruggedness of the aztreonam resistance landscape. (C) SHAP heatmap for the 553 most aztreonam-resistant genotypes (∼1% of the library). Compared to ampicillin selection, SHAP contributions exhibit greater heterogeneity, highlighting extensive epistatic interactions among mutations. (D) Beeswarm plot of SHAP values aggregated across all genotypes. While the absence of R244C/S mutations again confers significant positive effects, several other mutations (E240K, E104K, R164N/S/H, Q39K, A237T, and G238S) show context-dependent positive effects, emphasizing complex, combinatorial interactions. (E) Mean absolute SHAP values rank mutation positions by their overall influence on aztreonam resistance. Absence of mutations at position R244 remain most influential, but additional secondary positions emerge prominently, illustrating the multidimensional nature of the adaptive landscape. Together, panels A–E indicate that predicting aztreonam resistance demands larger training datasets and highlights intricate, context-dependent mutation interactions that differ significantly from those observed under ampicillin selection.

Although the LightGBM model predicted fitness effectively, it still functions largely as a “black box” and does not inherently reveal biological mechanisms such as epistasis. To address this, we interpreted the model using SHAP (SHapley Additive exPlanations) values, providing insights into how individual mutations contribute to predicted fitness and enabling visualization of epistatic interactions otherwise obscured by the high dimensionality of the data. In this context, SHAP values quantify the contribution of each specific mutation (e.g., Q39K, N276D) to the predicted fitness of a given genotype. A positive SHAP value indicates that the mutation increases predicted fitness relative to the average, whereas a negative value indicates a detrimental effect. Figure 6C summarizes SHAP scores for the top 553 most resistant variants (∼1% of the library) under ampicillin selection, allowing visualization of how the presence or absence of specific mutations contributes to AUC-Fitness. We further generated beeswarm plots (Figure 6D) to aggregate SHAP values across the dataset, identifying mutations with the greatest overall impact on fitness predictions. SHAP dependence plots revealed how the presence (red) or absence (blue) of each mutation influences its contribution to predicted fitness. Under ampicillin selection (Figure 6D), the most critical residue was R244, as variants with R244C and R244S mutations had lower AUC-Fitness, and fitness was generally higher when these mutations were absent. Mean absolute SHAP values (Figure 6E) further summarized the magnitude of each mutation’s influence on fitness.In contrast, the SHAP score distribution was more complex under aztreonam selection (Figure 7C). The heatmap of SHAP scores for the top 553 mutants showed a more convoluted pattern compared to ampicillin (Figure 7C). Although R244S/C again had very high absolute SHAP scores (Figure 7D–E), aztreonam resistance was generally higher when these mutations were absent, underscoring the importance of residue R244 for maintaining β-lactamase activity. Mutations including E240K, E104K, R164N, R164S, and R164H largely contributed positively to aztreonam resistance (Figure 7D–E). Interestingly, for G238S, both the presence and absence of the mutation sometimes increased resistance, suggesting complex and context-dependent epistatic interactions.

By examining these dependencies, we uncovered non-linear and epistatic effects critical for understanding the complex fitness landscape of antibiotic resistance evolution. Ultimately, combining machine learning predictions with SHAP interpretation not only enables accurate fitness prediction but also provides biologically interpretable hypotheses about how specific mutations and their interactions shape resistance. The generated SHAP values and visualizations offer a granular, interpretable view of the model’s internal logic, bridging the gap between predictive modeling and biological insight.

### Epistatic constraints revealed by evolutionary modeling define mutational paths toward ESBL activity in TEM-1 β-lactamase

To evaluate how well the TEM-1^CML^ sampled the underlying complete fitness landscape for TEM-1 from an evolutionary perspective, we analyzed pairwise and single-site statistics for the β-lactamase protein family (InterPro: PF13354) using **direct coupling analysis (DCA)**, a joint probability coevolutionary model that infers the family couplings (*e_ij_*) and local fields (*h_i_*) parameters of a Potts model of sequence composition^17,18^. While computing the family sequence logo, we found that although several amino acid alternatives were tolerated at the targeted sites within the broader family (**Figure 8A**), these sites are highly entrenched within the specific context of TEM-1^WT^ sequence. Conditional mutational probabilities computed from DCA parameters revealed that the effective alphabet (a limited number of available changes at a given site) at these positions in TEM-1 is highly restricted: with the exception of residues E104 and E240, all other targeted sites were largely limited to a single preferred residue (**Figure 8B**). This finding implies that, for most positions, only variants involving either mutations at 104 or 240—or coordinated, higher-order combinations of mutations—would be able to beneficially impact the epistatic sequence information of TEM-1, especially for maintaining TEM-1’s native substrate activity.

**Figure 8:**
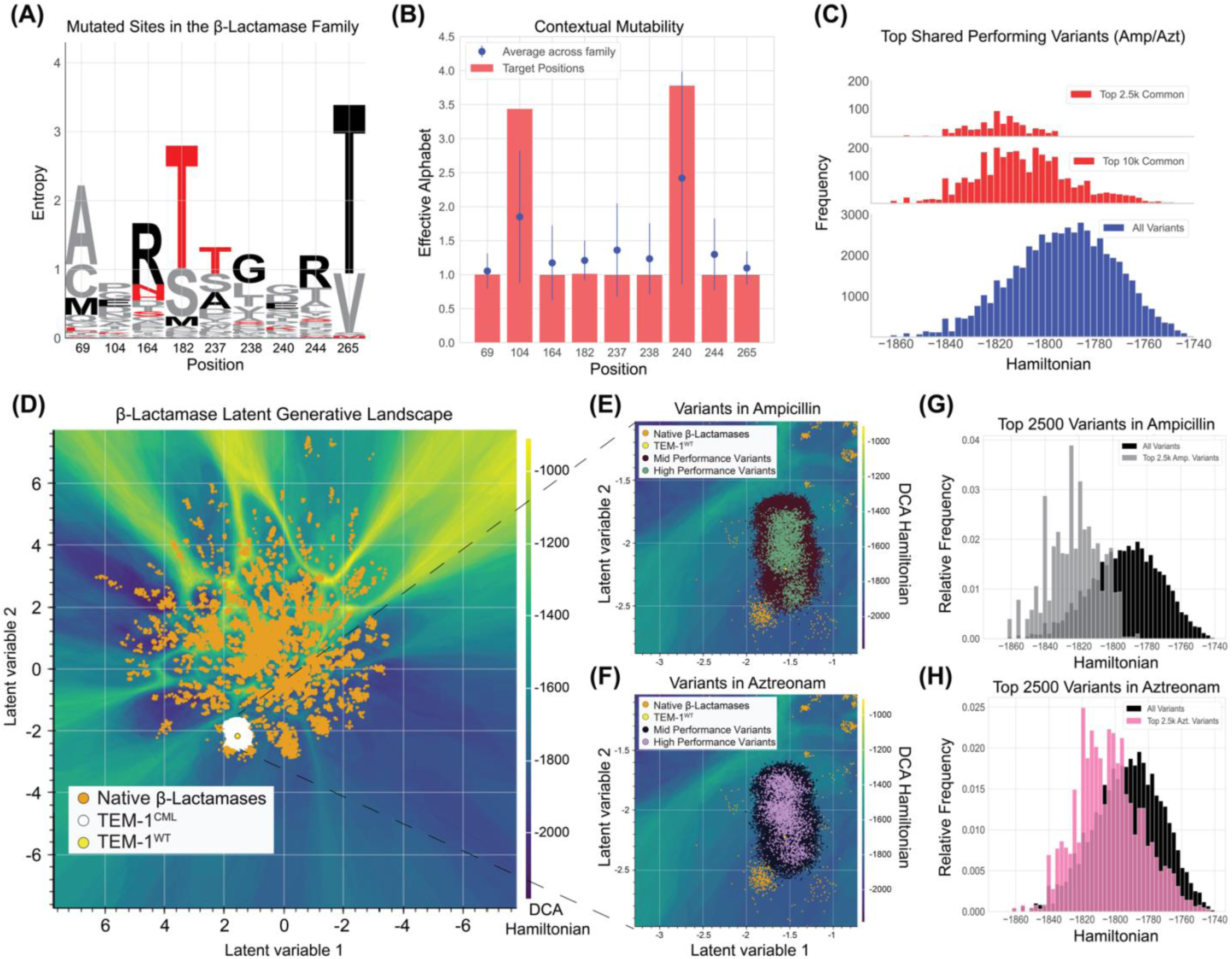
Evolutionary statistics of the TEM-1 family captures enzymatic optimization and epistatic requirements for ESBL in highly resistant variants. (A) β-lactamase family (PF13354) sequence logo for each site position in the mutational library. TEM-1 wildtype amino acids listed on x-axis. (B) Effective alphabet for each position based on the context mutability of the TEM-1 *E. coli* sequence as compared to the mutability of those sites for the whole family. (C) Hamiltonian distributions of the top 2,500 and 10,000 common performing variants (red) when exposed to both ampicillin ([AMP] = 781 µg/mL) and aztreonam ([AZT] = 36 µg/mL) compared against the distribution of all other variants. (D) Latent generative landscape for the β-lactamase family (PF13354) with native proteins labeled in orange. Regions of favorable Hamiltonian are colored dark blue and unfavorable regions (barriers) are shown in light yellow. The TEM-1^CML^ variants in white are located within the well that contains the TEM-1^WT^ sequence (yellow). (E) and (F) Zoomed in portion of the LGL where variants with increasing performance from both the ampicillin (E) and aztreonam (F) experiments ([AMP] = 781 µg/mL and [AZT] = 36 µg/mL) co-localize to the center of the well that contains the TEM-1^WT^ sequence. (G) and (H) The relative frequency of variants and the respective Hamiltonian energies for ampicillin (G) and aztreonam (H) experimental fitness scores for the top 2500 variants.

Using the DCA-derived couplings and fields, we computed a proxy for fitness—termed **sequence Hamiltonian**—based on the Hamiltonian of the DCA probability distribution (*Methods*) ^19,20^. In previous work, we validated the Hamiltonian as a metric for enzymatic activity and evolutionary fitness in TEM-1^21^. When analyzing the top 2500 variants with high AUC-Fitness under ampicillin selection (**Figure 8G**) or aztreonam (**Figure 8H**) selection, we observed that these high-performing variants were enriched for more negative (favorable) Hamiltonian scores compared to lower-fitness variants. A significant correlation between Hamiltonian scores and AUC-Fitness was present across different antibiotic concentrations (**Figure S6**). Moreover, this enrichment was apparent among variants that exhibited high fitness under both ampicillin and aztreonam selection (**Figure 8C**), suggesting that achieving extended-spectrum β-lactamase activity requires not only adaptations conferring broad substrate recognition but also compliance with evolutionary (and biophysical) constraints preserved within the β-lactamase family.

We further analyzed the β-lactamase family by creating a **latent generative landscape (LGL)** ^22^, a dimensionality reduction and generative method that maps protein sequences based on both functional properties and sequence Hamiltonian (**Figure 8D**). The LGL for β-lactamase family revealed that variants in our TEM-1^CML^ library clustered within the latent space surrounding TEM-1^WT^, with wells of favorable Hamiltonian scores (dark blue) and barriers of unfavorable sequence scores (light green). Both **Figure 8E** and **Figure 8F** demonstrate that the best-performing variants co-localized with wells of favorable Hamiltonian values, closely associated with the native TEM-1 sequence, under both ampicillin and aztreonam selection.

Collectively, these results support the hypothesis that retaining favorable epistatic interactions—as reflected by favorable Hamiltonians—is essential for preserving enzymatic function while acquiring resistance to new β-lactams. Although compliance with family-derived evolutionary constraints may not be sufficient alone for achieving extended-spectrum resistance—additional neofunctionalization events are likely required—it remains a critical evolutionary prerequisite that improves the probability of selecting functional, resistant variants.

## DISCUSSION

We engineered a comprehensive combinatorial mutant library (TEM-1^CML^) consisting of 55,296 barcoded variants covering all possible combinations of 18 clinically relevant mutations across 13 residues in TEM-1 β-lactamase. By measuring variant fitness under increasing concentrations of ampicillin and aztreonam at multiple time points, we generated one of the most extensive genotype-phenotype datasets (∼8 million fitness measurements) on antibiotic resistance. As expected, TEM-1 exhibited high activity against ampicillin and robustness to mutations, with most variants retaining measurable activity. Consistent with previous single-mutant studies, none of the tested combinations significantly exceeded wild-type TEM-1’s ampicillin resistance. The ampicillin fitness landscape proved highly predictable, as both linear regressions using single-mutant fitness values and machine-learning models trained on subsets of variants accurately reconstructed the landscape. Conversely, the fitness landscape under aztreonam selection was relatively more rugged, characterized by widespread high-order epistasis. We identified hundreds of aztreonam-resistant variants, demonstrating substantial evolvability of TEM-1 toward ESBL activity.

Graph-theoretic analysis revealed fundamental differences in evolutionary trajectories under the two antibiotics: ampicillin selection generated a landscape with few dominant peaks, whereas aztreonam selection fragmented the landscape into numerous local maxima exhibiting strong non-monotonic dependence on drug concentration. This variation in landscape topology underscores how epistasis and selection pressure shape evolutionary accessibility and diversity in resistance outcomes.

To further probe fitness predictability, we employed a LightGBM-based machine learning model coupled with SHAP values for interpretability. This approach provided accurate predictions for ampicillin resistance and moderate accuracy for aztreonam resistance. SHAP analysis highlighted both global and context-dependent mutational effects, identifying the pivotal role of residues A237 and R244 under ampicillin selection. In contrast, aztreonam resistance involved highly epistatic combinatorial effects from mutations such as E240K, E104K, and R164H/S/N. Intriguingly, mutations like G238S and Q39K demonstrated highly context-dependent effects, enhancing resistance in some genetic backgrounds while impairing it in others.

Additionally, we evaluated how effectively TEM-1^CML^ sampled the broader evolutionary sequence space using direct coupling analysis (DCA). DCA revealed strong evolutionary constraints at most targeted sites in TEM-1, except for residues E104 and E240. Consistent with these constraints, the majority of variants underperformed relative to wild-type TEM-1 under ampicillin selection. Experimental fitness correlated well with DCA-derived Hamiltonians under both antibiotics, and high-fitness variants consistently maintained favorable evolutionary couplings. Latent generative landscape (LGL) analysis further indicated that high-performing variants clustered near wild-type TEM-1 within optimal sequence-Hamiltonian regions, emphasizing the key importance of preserving epistatic compatibility during functional adaptation.

In summary, we present an integrated experimental and theoretical framework for studying β-lactamase evolution. By combining comprehensive mutant libraries, graph theory, interpretable machine learning (SHAP analysis), and evolutionary statistics, we provide a detailed understanding of the TEM-1 fitness landscape and evolutionary pathways leading to ESBL activity.

Our combined approach supports the rational design of mutation libraries to effectively explore functional sequence space for antibiotic resistance genes, enabling the development of testable hypotheses about the evolution of antibiotic resistance. Future work will apply this methodology to investigate the evolutionary dynamics of other clinically significant β-lactamases, such as the emergence of carbapenemase activity in cephalosporinases.

## ACKNOWLEDGEMENTS

E.T. is supported by UTSW Endowed Scholars Program, Welch Foundation I-2082-20240404, and NIGMS Grant (R01GM125748). FM, SA and JAP acknowledge support from the National Institute of General Medical Sciences grant R35GM133631. FM acknowledges support from NIH National Institute for Allergy and Infectious Diseases grant R01AI178692 and the National Science Foundation MCB-1943442.

## Methods

### Bacterial Strains

*Escherichia coli* NEB10-beta cultures were grown in Lysogenic Broth (LB) medium aerobically (shaking at 230 rpm) at 37°C, unless otherwise specified. Growth was determined by spectrophotometry (OD600). When specified, tetracycline was supplemented at 12 μg/mL. Three strains were used in this study: *E. coli* strain without plasmid, *E. coli* harboring pBR322-TEM-1, and *E. coli* containing pBR322-TEM-1^CML^. Plasmid-containing strains were maintained under tetracycline selection to ensure plasmid retention throughout all experiments.

### pBR322-TEM-1-S70A-E166A-Barcode Construction

The catalytically inactive TEM-1 mutant was generated using the pBR322-*blaTEM-1* plasmid as template. Site-directed mutagenesis to introduce the S70A and E166A mutations was performed via PCR amplification using Q5 High-Fidelity DNA Polymerase (New England Biolabs). The PCR products were assembled using Gibson Assembly Master Mix (New England Biolabs) according to the manufacturer’s instructions. The assembled plasmid was transformed into electrocompetent Escherichia coli NEB10-beta cells following standard electroporation protocols. Plasmid DNA was extracted and purified using NucleoSpin Plasmid Miniprep Kit (Macherey-Nagel), and the presence of the desired mutations was confirmed by sequencing (Plasmidsaurus, USA). Next, a unique barcode sequence was introduced downstream of the mutated *blaTEM-1* gene using Gibson Assembly. The barcoded plasmid construct was transformed into NEB10-beta cells, and several colonies were selected for sequencing to confirm the presence of the barcode sequence.

### pBR322-TEM-1-Wild-type-Barcode Construction

For the barcoded wild-type construct, the pBR322-*blaTEM-1* plasmid was used as a template. A unique barcode sequence was introduced downstream of the wild-type *blaTEM-1* gene using Gibson Assembly with the appropriate primers. The barcoded construct was transformed into *E. coli* NEB10-beta cells by electroporation according to the manufacturer’s protocol. Single colonies were isolated and sequenced to confirm the successful incorporation of the barcode.

### Combinatorial Mutant Library Preparation

The combinatorial mutant library consisting of 55,296 variants was synthesized by Twist Bioscience on a low-copy number plasmid, pBR322, to better replicate environmental conditions. Each variant in the library included a unique 52-base pair barcode downstream of the gene, containing 15 strategically placed positions with four different possible nucleotides (A, T, G, C) at each position. This design created a diverse set of possible sequences, enabling the generation of up to 10^9 unique barcodes for precise mutant identification. For library transformation, 100 ng of the combinatorial mutagenesis plasmid was electroporated into 25 μL of *NEB10-beta E. coli* cells in ten separate replicates. After electroporation, each replicate was immediately mixed with 975 μL of NEB10-beta/stable outgrowth medium and incubated at 37°C with shaking for 1 hour. To assess transformation efficiency, 10 μL from each outgrowth culture was removed for analysis. The remaining 990 μL was transferred to 20 mL of LB medium supplemented with tetracycline (12 μg/mL) and cultured overnight at 37°C with shaking. Nine of the ten replicates achieved transformation efficiencies exceeding 10^8 transformants. These successful replicates were pooled at equivalent OD600 ratios, divided into glycerol stocks, and stored at -80°C until used for experiments.

### Growth Assay for Fitness Measurements

*E. coli* + pBR322-TEM-1-Barcode and catalytically inactive *E. coli* + pBR322-TEM1-S70A-E166A-Barcode strains were inoculated in Luria-Bertani (LB) broth supplemented with tetracycline (12 μg/mL) and incubated overnight at 37°C with shaking at 230 rpm. In parallel, three replicate samples of the combinatorial mutant library were thawed and individually transferred to Erlenmeyer flasks containing 100 mL of LB broth supplemented with tetracycline (12 μg/mL), then incubated under identical conditions. The following day, all cultures were diluted to an optical density (OD600) of 0.02 in fresh LB broth containing tetracycline and grown for 1.5 hours at 37°C with shaking. After this initial growth period, each replica of the combinatorial mutant library was spiked with 5% (v/v) of the TEM-1-Barcoded cell line and 1% (v/v) of the TEM-1-S70A-E166A-Barcoded cell line. The mixed cultures were then diluted to an OD600 of 0.01. 24 mL of the culture were transferred to 50 mL conical tubes for testing across different antibiotic concentrations. Aztreonam was tested at the following concentrations: 3000, 324, 108, 36, 12, 4, 1.33, 0.44 μg/mL, and a no-drug control. For the ampicillin assay, cultures were exposed to: 50000, 12500, 3125, 781, 195, 48.8, 12.2, 3.1 μg/mL, and a no-drug control. Optical density (OD600) was measured every 3 hours up to 12 hours. At each concentration and time point, 5 mL of culture was collected, centrifuged for 5 minutes at 6,000g, and the cell pellet was stored at -20°C for subsequent plasmid isolation.

### NGS Sample Preparation

Pellets obtained from the Competitive Growth Assay were thawed, and plasmids were isolated using the NucleoSpin Plasmid isolation kit (Macherey-Nagel) following the manufacturer’s instructions. Plasmids were eluted in 35 μL of water, and concentrations were assessed using the Qubit Assay. Samples that yielded sufficient plasmid concentration were used for further sample preparation. The 52 bp barcode region, located 30 bp downstream of the blaTEM-1 gene, was amplified using primers positioned 75 bp upstream (forward primer: 5’-GACAGATCGCTGAGATAGGTGCCTC-3’) and 80 bp downstream of the barcode (reverse primer: 5’-CTAGGAAAAACTATTAGAGTACTGGTTTTAGGG-3’), generating a 207 bp PCR product. PCR cycling conditions consisted of initial denaturation at 95°C for 3 minutes, followed by 25 cycles of: 95°C for 15 seconds, 65°C for 20 seconds, and 72°C for 30 seconds. A final extension step was performed at 72°C for 3 minutes. The amplified region was purified using NucleoSpin Gel and PCR Clean-Up (Macherey-Nagel) according to the manufacturer’s protocol. The concentration of purified PCR products was assessed using the Qubit assay. Samples containing at least 1000 ng of DNA in a 50 μL volume were submitted for NovaSeq X Plus 300 cycle sequencing.

### Minimum Inhibitory Concentration (MIC) Measurements

*E. coli* strains harboring either no plasmid, pBR322-TEM-1, or pBR322-TEM-1CML were cultured overnight in 5 mL Luria-Bertani (LB) medium at 37°C with shaking at 230 rpm. Tetracycline (12 μg/mL) was added to the media for strains containing plasmids, while the plasmid-free NEB10-beta strain was cultured in plain media. The next day, cultures were diluted to an optical density (OD600) of 0.001 in fresh medium. The diluted cultures were exposed to 3-fold serial dilutions of various β-lactam antibiotics (ampicillin, ceftazidime, cefotaxime, ceftriaxone, aztreonam, cefepime, ertapenem, imipenem, meropenem) in 96-well plates. Plates were incubated at 37°C with shaking (400 rpm in Infors HT plate shaker). Minimum inhibitory concentration (MIC) values were determined by measuring optical density using a Biotek Epoch2 plate reader at the 20h timepoint. All experiments were performed in triplicate. For each strain and tested antibiotic, MIC was defined as the lowest concentration where growth was below 5% of the maximum growth observed in the dose-response curve. Background correction was applied to all OD measurements using the median background from control wells. For biological replicates, MIC values were calculated independently for each replicate and then averaged. Statistical analysis of MIC values between strains was performed using two-tailed unpaired t-tests with unequal variance (Welch’s t-test). Significance levels were indicated as follows: * for p < 0.05, ** for p < 0.01, and *** for p < 0.001.

#### Fitness Landscape Graph Construction

The protein fitness landscape was encoded as a directed, weighted graph *G* = (V, E), with each node 𝑣 ∈ *V* representing a unique mutant and each edge (*u*, 𝑣) ∈ *E* connecting variants that differ by exactly one amino acid substitution. Edge orientation follows increasing fitness—pointing from *u* to 𝑣 if *f*(𝑣) > *f*(*u*), and bidirectional when *f*(𝑣) = *f*(*u*)—and weights are defined as 𝑤(*u*, 𝑣) = exp|*f*(*u*) − *f*(𝑣)|, so that log-scale fitness differences are exponentially accentuated. Although *G* captures every single-mutation path, its dense connectivity precludes meaningful visualization or interpretation. Therefore, we merge nodes into supernodes containing collections of mutants that can convert to each other via neutral point mutations as follows.

We first define an edge to be a neutral edge if 𝑤(*u*, 𝑣) ≤ 𝑒^𝜃neutral^. We determined the neutral threshold by plotting the distribution of fitness differences between all pairs of mutants that differ by a single amino acid substitution under no drug conditions, and choosing 𝑒^𝜃neutral^ to be the fitness difference amplitude below which 99% of all such pairs differ. We apply a neutral merging algorithm that combines nodes that can convert to each other via neutral edges into supernodes. This merging is implemented with a Union-Find data structure for near-linear time complexity. In each iteration, after merging neutral (super)nodes into the next iteration of supernodes, we denote the partition of the single-mutant nodes into the current iteration *t* of supernodes {S_i_(t)} (initially {S_i_(0)} = {v_i_}). We define the net flow edge weight between two supernodes 𝑆_*i*_(*t*) and 𝑆_𝑗_(*t*) to be the aggregated value of all edges between mutants that connect the two supernodes: 𝑤_net_(𝑆_*i*_(*t*), 𝑆_𝑗_ (*t*)) = |∑_*u*∈𝑆_*i*_(*t*), 𝑣∈𝑆_*j*_(*t*)_ 𝑤(*u*, 𝑣) − ∑_*u*∈𝑆_*i*_(*t*), 𝑣∈𝑆_*i*_(*t*)_ 𝑤(*u*, 𝑣)|. Two nodes from the previous iteration 𝑆_*i*_(*t* − 1), 𝑆_𝑗_ (*t* − 1) can be merged into a single node if there exist *u* ∈ 𝑆_*i*_ (*t* − 1),𝑣 ∈ 𝑆_𝑗_ (*t* − 1) such that 𝑤(*u*, 𝑣) ≤ 𝑒^𝜃neutral^, unless there exist *u* ∈ 𝑆_*i*_ (*t* − 1),𝑣 ∈ 𝑆_𝑗_(*t* − 1) such that (*u*, 𝑣) ∈ 𝐹, where *F* is the set of forbidden pairs (see below for determination of forbidden pairs). After each merge, we recompute the net-flow weights to all other supernodes as above, and iterate until no further merges are allowed.

Because a single path of neutral edges can connect two large supernodes that are otherwise connected by edges with large fitness differences—thereby obscuring the dominant feature, we first identified a set of forbidden pairs of supernodes, which cannot be merged in our algorithm. To identify the forbidden pairs, we first applied a conservative “tiny” neutral merge on the original graph *G*. Any two nodes joined by an edge with w ≤ 𝑒^θtiny^ (𝜃_tiny_ = 0.02) were collapsed into a single supernode. From this preliminary graph, we then identified the set *F* of *forbidden pairs*—edges whose weights exceed 𝑒^θlarge^. Formally, 𝐹 = {(*u*, 𝑣) ∈ *E* ∣ 𝑤(*u*, 𝑣) ≥ 𝑒^𝜃large^}, with 𝜃_large_ = 5.5 chosen to exceed any realistic single-mutation effect. Only the forbidden pair with largest edge weight is prevented to be merged in the following process.

From the resulting coarse-grained fitness landscape graph, we annotated peak nodes (nodes with in-degree > 0 and out-degree = 0), connection nodes (variants that lie on evolutionary routes to two or more peaks, i.e. those in the ancestor sets of multiple peaks), and peak committed edges (those leading exclusively toward a single peak’s ancestor set).

### Machine Learning Analysis of TEM-1 β-lactamase Genotype-Fitness Relationship for AZT and AMP Resistance

This analysis investigates how mutations in the TEM-1 β-lactamase enzyme influence its functional fitness regarding resistance to Aztreonam (AZT) and Ampicillin (AMP). We employ a LightGBM regression model to build a predictive genotype-phenotype map, capturing complex, non-linear interactions among mutations. Model performance is validated through learning curves and evaluation on held-out test data to ensure predictive capability.

To interpret this “black box” model, we utilize SHAP (SHapley Additive exPlanations) values, grounded in cooperative game theory. SHAP values assign a quantitative importance to each mutation (e.g., L21P, N276D), measuring how each mutation shifts predicted fitness from the baseline (the average fitness across all mutants). Positive SHAP values indicate mutations driving predictions higher (enhancing AZT or AMP resistance), while negative values indicate reductions in predicted fitness.

The robustness of SHAP interpretation rests on key theoretical properties: (i) Local Accuracy ensures feature contributions sum exactly to the difference from baseline for each prediction; (ii) Missingness assigns zero importance to absent mutations; and (iii) Consistency guarantees SHAP values do not decrease if a mutation’s actual influence increases or stays constant upon model retraining.

Through global summary (beeswarm) plots, SHAP values identify mutations with significant overall impacts, highlighting key resistance determinants. Additionally, SHAP dependence plots reveal detailed insights into individual mutation effects and epistatic interactions, illustrating how one mutation’s effect can depend on the presence or absence of others. Combining predictive modeling with SHAP interpretation provides a biologically meaningful approach, generating testable hypotheses on mutation impacts and their combinations, thus bridging predictive accuracy and functional understanding of antibiotic resistance evolution.

### Direct Coupling Analysis on **β**-lactamase enzyme family

We used HMMSearch^23^ against the Uniprot database (Swiss-Prot and TrEMBL) to obtain MSA for the protein family β-lactamase (PF13354) using the PFAM HMM seed^24^ The original MSA (104,744 sequences) was filtered to remove sequences with 5% or greater contiguous gaps (27,242 sequences after filtering). We then applied the DCA method^17^ to the family MSA to estimate the direct coupling between all pairs of residues, as well as the residue preferences at each position. DCA utilizes maximum entropy modeling to postulate a Potts-like joint probability distribution of protein sequences

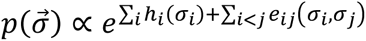

where the position of residues within the aligned domain or protein sequence is denoted as i and j, and parameters e_ij_ and h_i_ are numerically inferred via a mean field python implementation [https://github.com/utdal/py-mfdca]. The e_ij_ parameters quantify the coupling strength for residues i and j for all possible amino acid occurrence pairs. The amino acid preferences for independent positions are captured by the parameter h_i_.

Both contributions from couplings are captured by the negative log-likelihood of the model, commonly referred to as Hamiltonian (H):

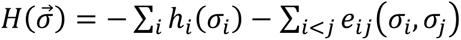

The Hamiltonian score quantifies global similarity to the training alignment and has been used as a proxy measure of sequence fitness^19,20^. To assign a Hamiltonian score to the sequences at TEM-1^CML^, we took the overlapping region of TEM-1 with the Pfam (PF13354) domain (position 48 to 261 for Uniprot: P62593), leaving out positions 21, 39, 275, and 276 from the 13 mutated positions.

### Context-dependent Entropy and Effective Alphabet

Given the probability distribution given by DCA, it is possible to quantify how restricted a position is given the rest of the sequence by focusing on the conditional probability of the site:

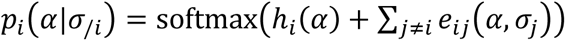

By computing the statistical entropy associated with this distribution 𝑠_*i*_(𝜎⃗) = − ∑_*i*_ 𝑝_*i*_(𝛼|𝜎_/*i*_) log 𝑝_*i*_(𝛼|𝜎_/*i*_) we obtain a metric on the site restriction given its sequence context and the family parameters. Further, we can calculate an effective alphabet size, as described in^20^, for a more intuitive metric. The effective alphabet per site is given by

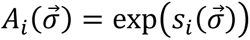

The effective alphabet ranges from one whenever a site is restricted to a single amino acid up to 21 when it is completely free to undergo substitutions to any of the 20 amino acids or a gap. This quantity does not depend on the current amino acid in site i, but on the state of the rest of the sequence; this allows us to directly quantify how the ensemble of correlated sites in the protein restricts the amino acid identity at a given position

### VAE Model Architecture and Latent Generative Landscapes

The Variational Auto-Encoder (VAE) is an unsupervised generative machine learning architecture that compresses high-dimensional data into a reduced space of latent variables (z). The architecture consists of three parts: 1) encoder network 𝑞_𝜙_(𝑧|𝜎⃗), 2) latent space representation 𝑧 = (𝑧_1_, 𝑧_2_), and 3) decoder network 𝑝_𝜃_(𝜎̂|𝑧). The encoder network maps the one-hot encoded input sequence 𝜎⃗ into a distribution over the bidimensional latent space 𝑞_𝜙_(𝑧|𝜎⃗). The latent variable z is a non-trivial representation of the essential features of the amino acid sequence with added noise that represents variations across the training dataset. Finally, the decoder associates a sequence 𝜎̂ with every point of the latent space; a more detailed description of the underlying architecture can be found in ^22^. The network is trained on the family MSA using the Adam optimizer to minimize the evidence lower bound function (ELBO):

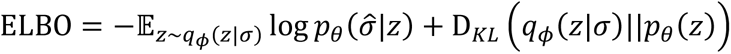

The first term is the reconstruction error, which measures how well the encoded data matches the generated data, while the second term serves as regularization to enforce a Gaussian shape of the conditional distribution over latent space for the encoder.

Once the VAE model has been trained, we take a grid over the latent space with uniform spacing and decode the corresponding sequences. Once decoded, we calculate the Hamiltonian score as described above and associate its value with the sampled point, i.e.: 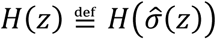.

---

**Figure S1.**
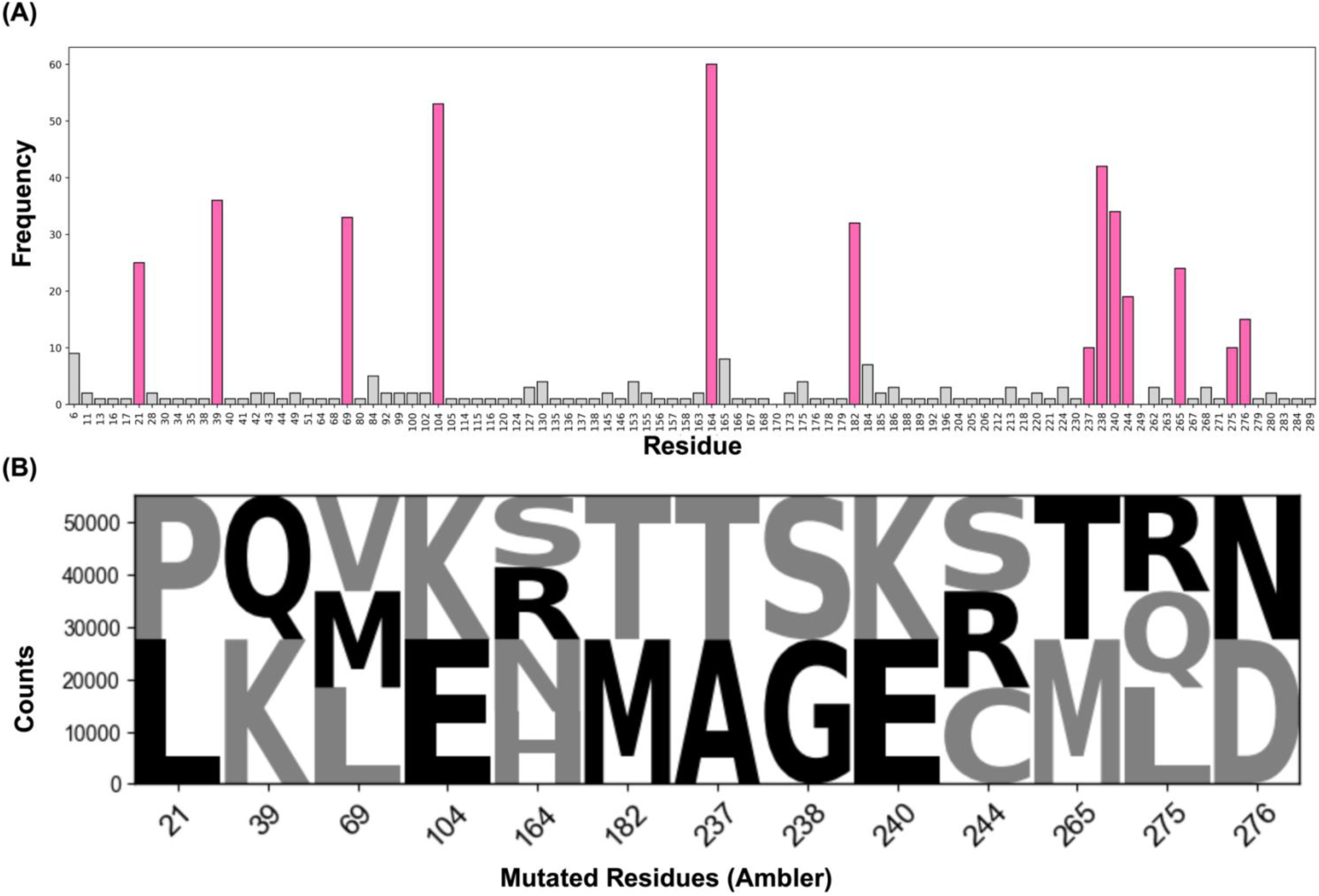

**Figure S2.**
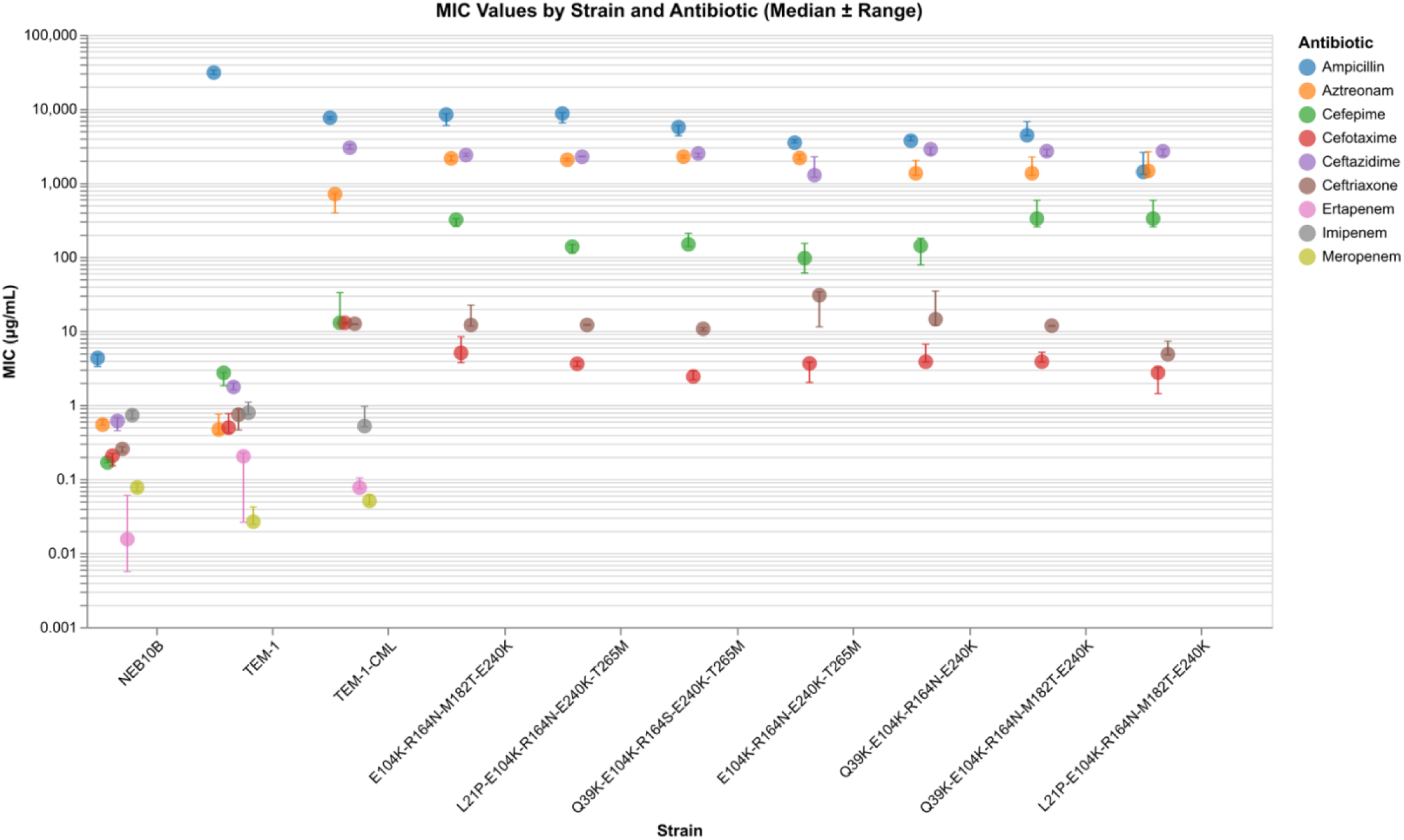

**Figure S3.**
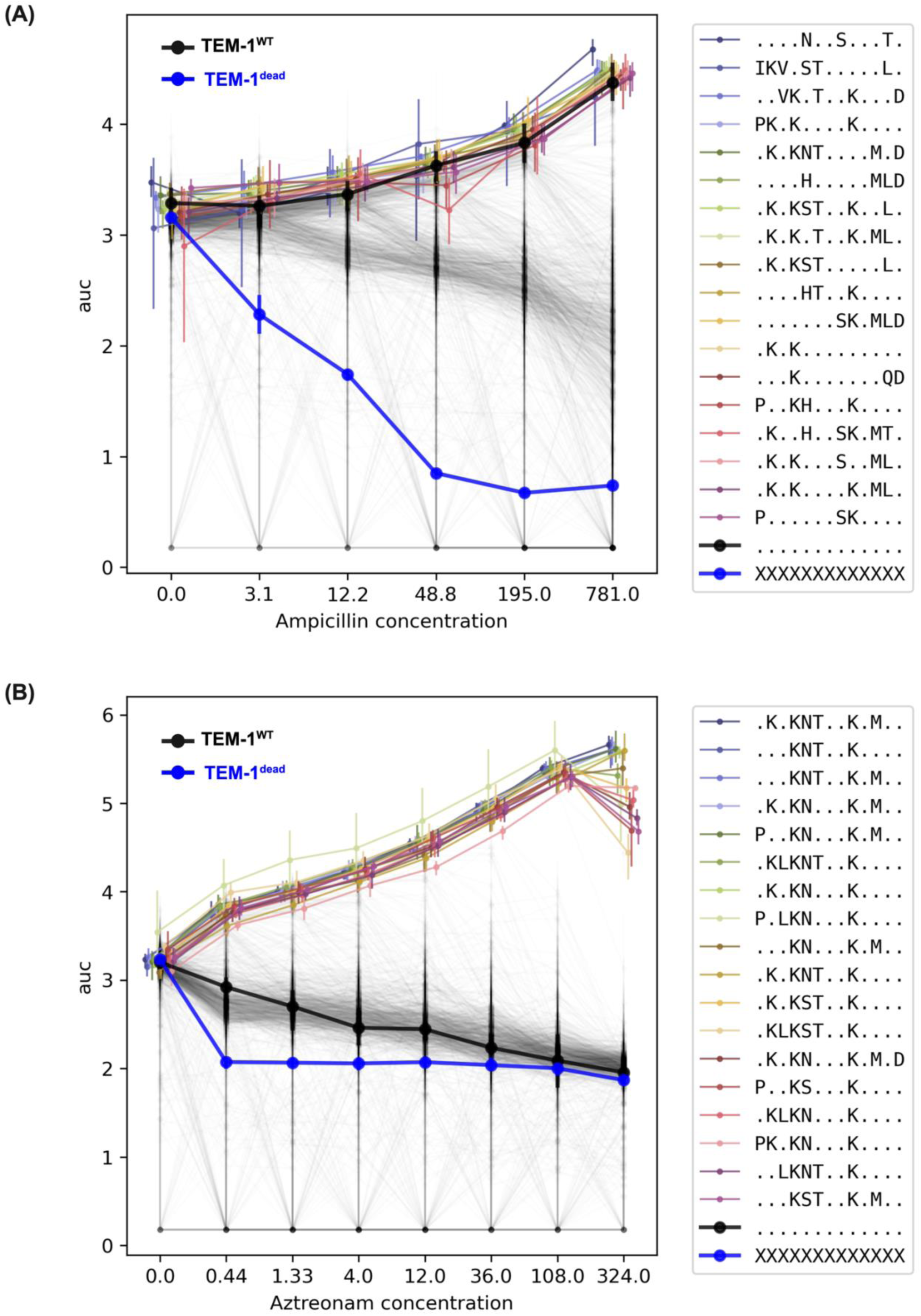

**Figure S4.**
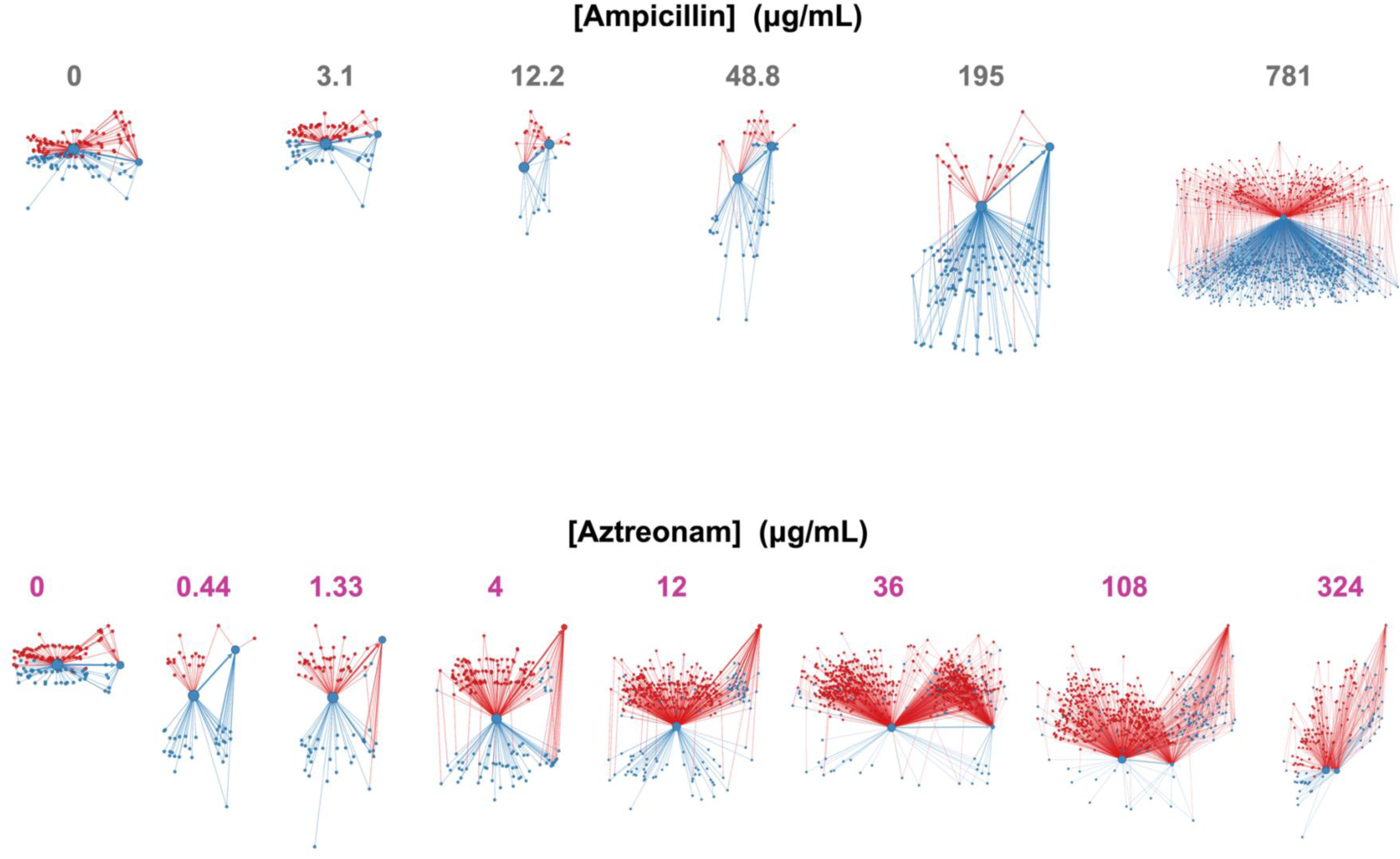

**Figure S5.**
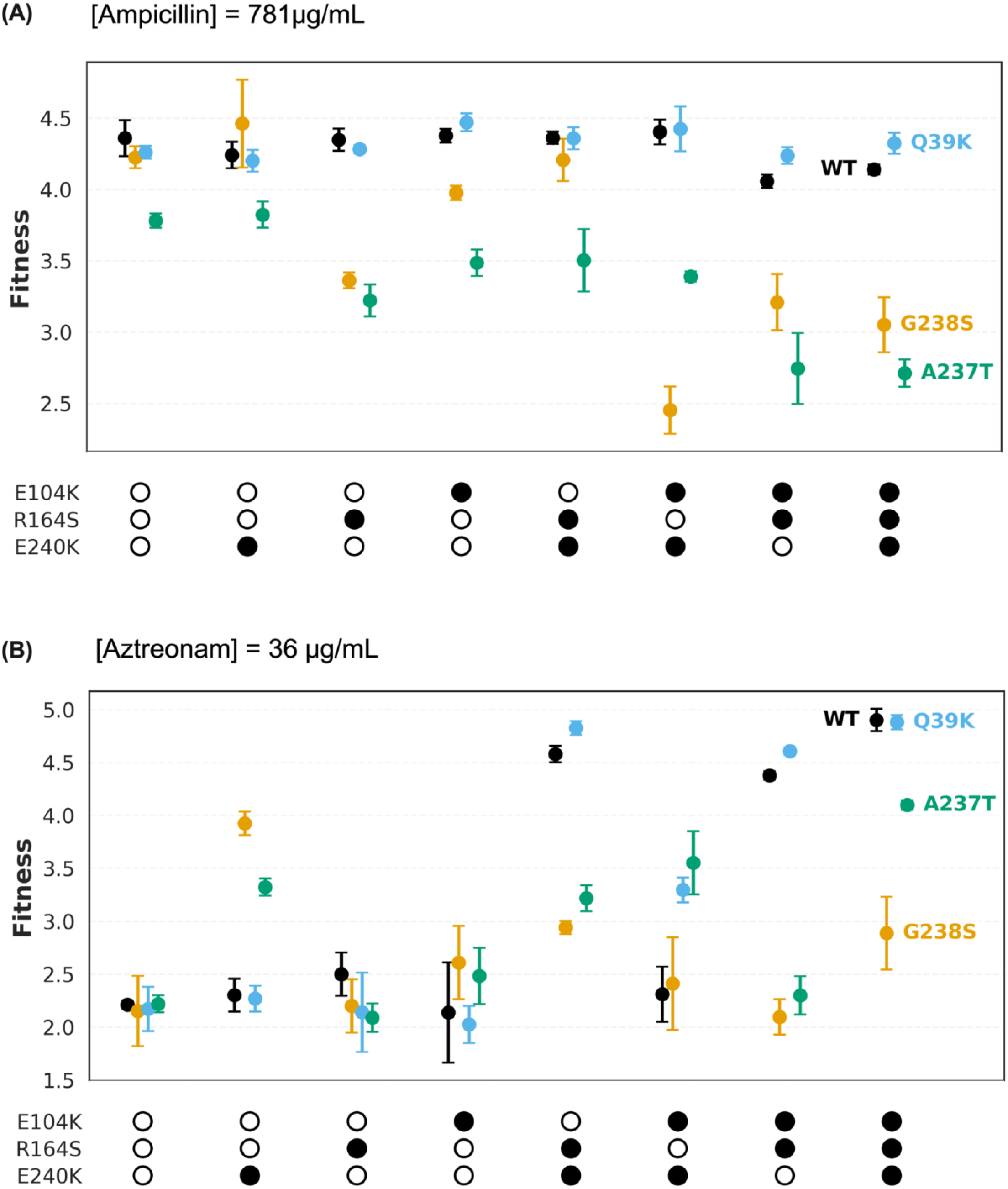

**Figure S6.**
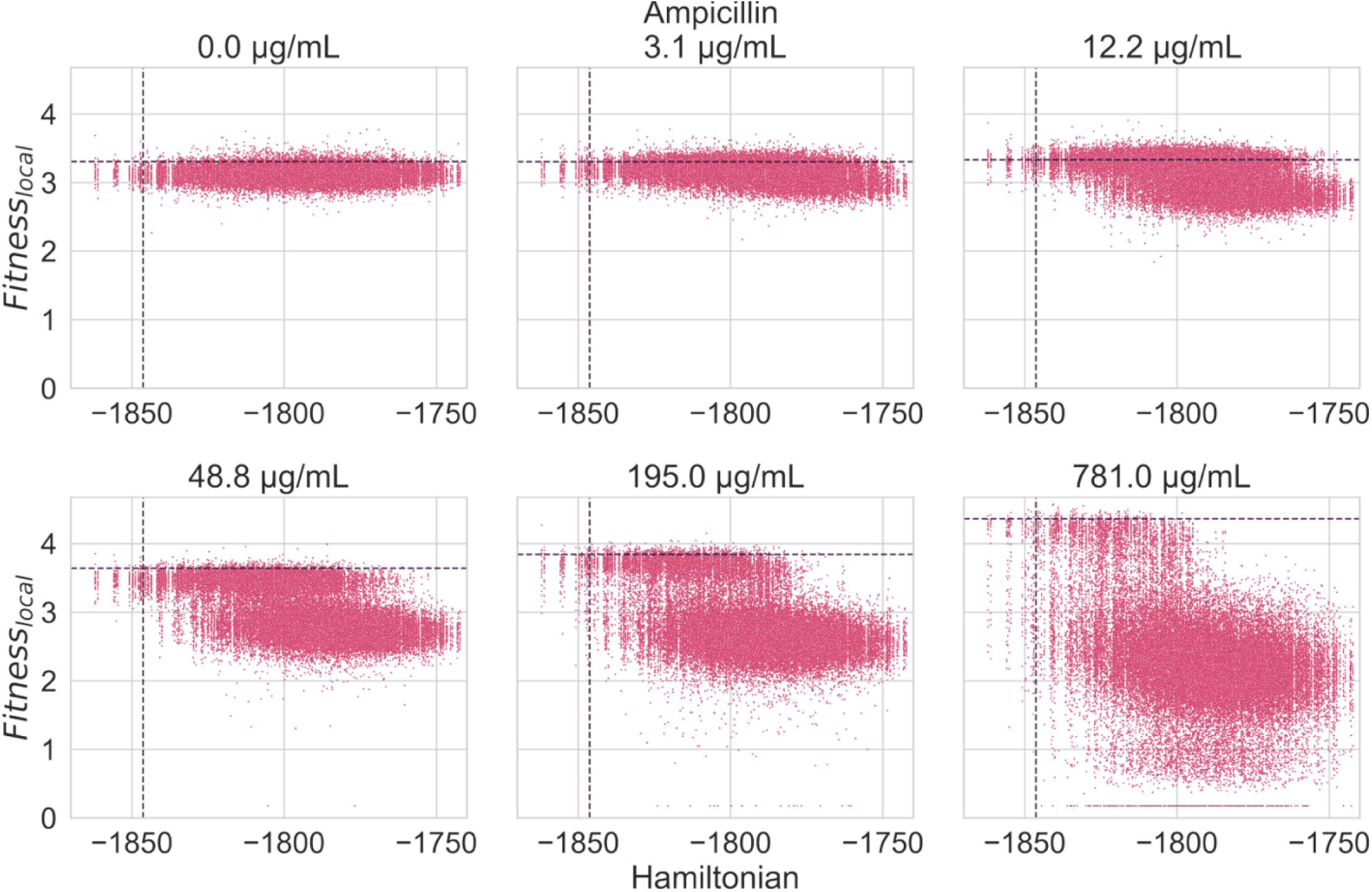
Analysis of TEM-1^CML^ fitness scores with evolutionary statistics. Correlation between fitness score and Hamiltonian energies at every concentration for Ampicillin. The vertical and horizontal dashed lines correspond to the Hamiltonian and AUC-fitness score for the TEM-1^WT^ sequence.

